# Structural characterization of the oligomerization of full-length Hantaan virus polymerase into symmetric dimers and hexamers

**DOI:** 10.1101/2023.10.04.560883

**Authors:** Quentin Durieux Trouilleton, Dominique Housset, Benoît Arragain, Hélène Malet

**Affiliations:** Univ. Grenoble Alpes, CNRS, CEA, IBS, F-38000 Grenoble; European Molecular Biology Laboratory (EMBL), Grenoble, France; Institut Universitaire de France (IUF)

## Abstract

Hantaan virus is a dangerous human pathogen whose segmented negative-stranded RNA genome is replicated and transcribed by a virally-encoded multi-functional polymerase. Here we describe the complete cryo-electron microscopy structure of Hantaan virus polymerase in several oligomeric forms. Apo polymerase protomers can adopt two drastically different conformations, which assemble into two distinct homodimers, that can themselves gather to form hexamers.

Polymerase dimerization induces the stabilization of most polymerase domains, including the C-terminal region that notably contains a C-terminal domain that contribute the most to dimer’s interface, along with a lariat region that participates to the polymerase steadying.

Binding to viral RNA induces significant conformational changes resulting in oligomer disruption, suggesting the possible involvement of multimers as protecting systems that would stabilize the otherwise flexible C-terminal domains.

Overall, these results provide new insights into the multimerization capability of Hantavirus polymerase and may help to define antiviral compounds to counteract these life-threatening viruses.

## INTRODUCTION

*Bunyavirales* is a large order of segmented negative stranded RNA viruses (sNSV) that encompasses several highly pathogenic zoonotic viruses^1^. Most Bunyaviruses are transmitted by mosquitoes or ticks, except Hantaviruses that are rodent-borne^2^. Human infection by Hantaviruses can result in severe diseases such as haemorrhagic fever with renal syndrome in the case of old-world Hantaviruses or encephalitis in the case of new-world hantaviruses^3^.

Representative members of the old-world hantaviruses are Hantaan and Puumala viruses whose infections result in fatalities in up to 15% of the cases^4^, whereas new-world Andes and Sin Nombre (SNV) hantavirus infections have mortality rates up to 40%^5^. Despite their potential threat for human health, neither drugs nor vaccines are presently available to combat or prevent infection by these types of viruses.

In this context, we focused our research on Hantaan virus polymerase (HTNV-L) that could be a key target for antiviral drugs development. It catalyses two fundamental steps of the viral cycle: replication that results in the duplication of the viral genome and transcription that generates viral messenger RNA (mRNA). As for other Bunyaviruses, the replication process uses an internal prime-and-realign mechanism for its initiation in the absence of a primer^6–8^. Conversely, transcription is performed by host-cell mRNA cap-snatching. This intricate process involves the binding of host-cell mRNA through the polymerase cap-binding domain (CBD) and their subsequent cleavage after 10 to 16 nucleotides by HTNV-L endonuclease (ENDO), thus generating capped RNA primers used by the polymerase core to elongate the viral mRNA^6,7^. Knowledge on the *Bunyavirales* polymerase organization comes from several structures that have been determined in the last few years. These include the structures of La Crosse virus (LACV, *Peribunyaviridae* family)^9–11^, Lassa and Machupo viruses (LASV and MACV, *Arenaviridae* family)^12–15^, Dabie Bandavirus (DBV, previously named Severe Fever with Thrombocytopenia Disease virus, *Phenuiviridae* family), and Rift Valley Fever virus (RVFV, *Phenuiviridae* family). In the case of HTNV-L, its first structural characterization came from the isolated ENDO domain that was determined by X-ray crystallography^16^. More recently, cryo-electron microscopy structures of HTNV-L and SNV-L have unveiled the organisation of Hantavirus polymerase core ^8,17^. The interior of the cores adopts a right-hand organization that is common to all RNA-dependent RNA polymerases (RdRp) and comprises a palm, a finger and a thumb domains. This central part of the core is encircled by (i) the linker region that connects the ENDO to the core, (ii) the core lobe that contains a vRNA-binding lobe (vRBL), (iii) the thumb-ring that surrounds the thumb and (iv) the lid that closes the active site cavity. The active site cavity is located in the core center at the intersection of four tunnels: the RNA template entry, the nucleotide entry, the template exit and the product exit tunnels^7,9^. The active site cavity contains the canonical catalytic motifs A to F that are conserved in all RdRp^18^. HTNV-L and SNV-L core structures were solved in an inactive configuration, wherein several of the catalytic motifs were found to be misplaced. In particular, the motif E, also known as “primer-gripper” which typically adopts a canonical 3-stranded β-sheet conformation, was shown to display an unusual α-helical conformation in the absence of RNA. Binding of the 5′vRNA end as a hook in a specific binding-site on the HTNV-L core was shown to induce a conformational rearrangement of the motif E into the more standard 3-stranded β-sheet structure, inducing several reconfigurations, notably the ordering of the putative prime-and-realign loop (PR loop) that might be involved in prime-and-realign initiation. Binding to the 5′vRNA end also induces the organization of motif F that is disordered in the apo form. In both HTNV-L and SNV-L structures, only the cores are visible as flexibility prevents the visualization of the ENDO and of all the C-terminal regions.

In this article, we present near-atomic resolution cryo-EM structures of the complete HTNV-L in various oligomeric states. The structures unveil the location of every domain of this central enzyme, notably depicting the organization of its C-terminal region that was uncharacterized. The astonishing flexibility of the polymerase is revealed, which results in the presence of two very distinct conformations of protomers, that can assemble into two singular homo-dimers. Dimerization implies a domain-swapping mechanism that could be conserved amongst Bunyaviruses. Our results in addition reveal that the homo-dimers can further assemble into a hexamer composed of a trimer of dimers, forming a unique intricate organization that had never been observed in any polymerases. Finally, we show that vRNA binding triggers large conformational changes in the protomers leading to multimer disruption and monomer activation, suggesting a role for multimers as possible storage systems that would stabilize and protect HTNV-L, prior to its activation for genome replication and transcription.

## RESULTS

### Structure of full-length HTNV-L monomer, dimer and hexamer

A full-length wild-type (WT) HTNV-L construct comprising an N-terminal his-tag was expressed in Hi5 insect cells and subsequently purified to homogeneity (**Supplementary** Fig. 1a and b). Through a combination of size-exclusion chromatography and mass photometry analysis, the presence of distinct HTNV-L oligomers was observed. These species encompass monomers (64%), dimers (17%) and a smaller fraction of higher molecular weight oligomers up to hexamers (5%) (**Fig. 1a**). Mass photometry also reveals the presence of a specie around 150kDa that could potentially correspond to the HTNV-L core only, too minoritarian to be detected in SDS-PAGE.

**Figure 1.**
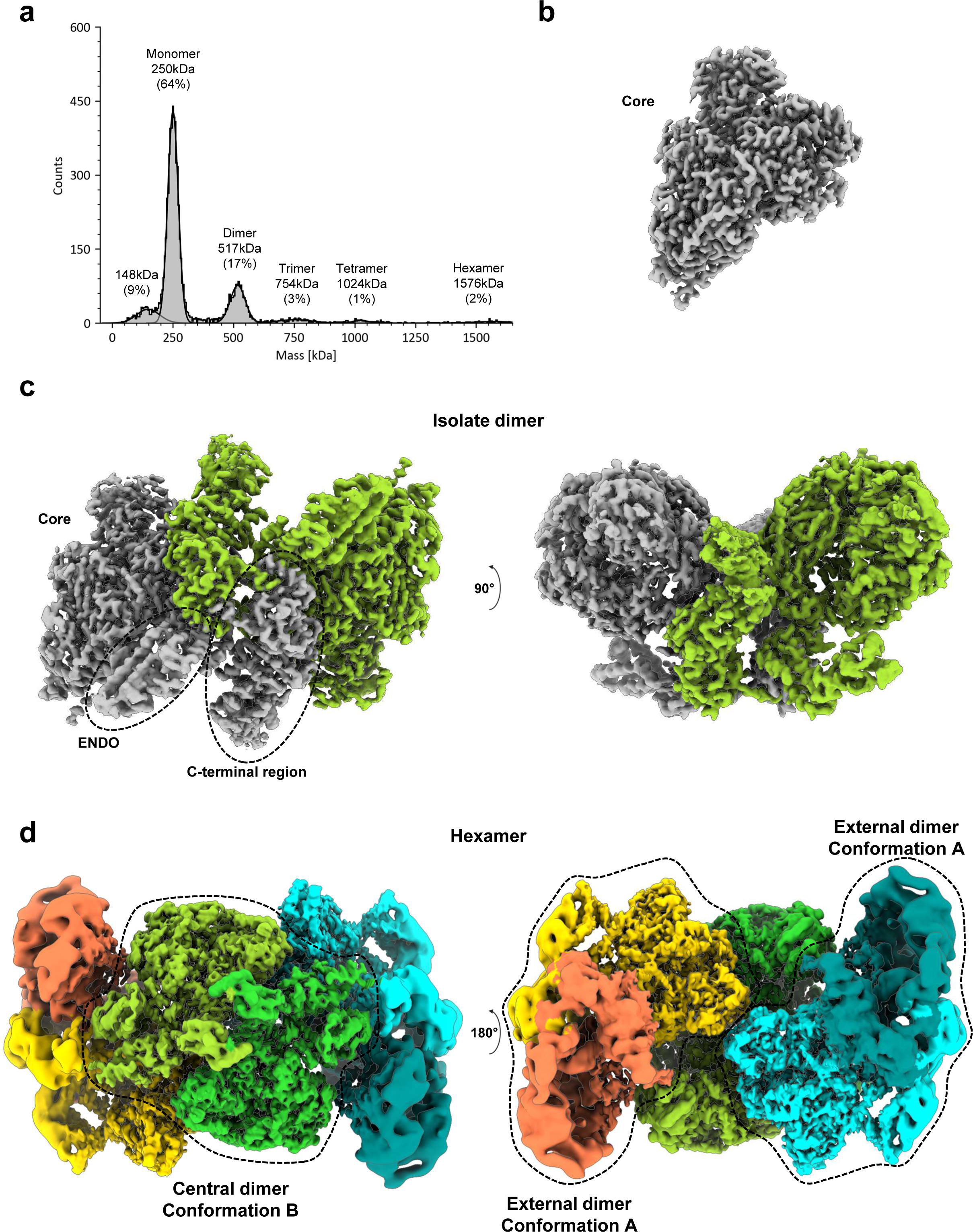
Oligomeric states of HTNV-L. **a** Mass photometry of HTNV-L. The gel filtration peak that corresponds to a 250kDa peak is analyzed. The molecular weight and the percentage of each specie is indicated. **b**, **c** and **d** Composite cryo-EM maps of HTNV-L in monomeric state (**a**), dimeric state (**b**) and hexameric state (**c**). Each protomer is colored differently. Dimers are labeled according to their respective conformations, named A and B.

The equilibrium between the various HTNV-L oligomers hinders any further biochemical separation (**Supplementary** Fig. 1c). Consequently, after cryo-EM data collection and initial image processing, particle picking was performed using both template and neural-network based picker^19,20^, thereby facilitating the selection and separation of monomers from both symmetric dimers and hexamers (**Supplementary** Fig. 2 and 3). From this dataset, multiple structures of apo HTNV-L were obtained in different oligomeric states:

i. The structure of monomeric apo HTNV-L WT, determined at an overall resolution of 2.6 Å, exhibits a conserved architecture compared to the previously obtained apo HTNV-L_D97A_^8^ (**Fig. 1b, Supplementary** Fig. 2**, Supplementary Table 1**). The polymerase core is clearly visible whereas the ENDO and the C-terminal region are too flexible to be resolved.
ii. The structure of an isolated apo HTNV-L WT symmetric dimer, that is determined at an overall resolution of 3.0 Å resolution, reveals the location of all domains, including the ENDO and the C-terminal region (**Fig. 1c, Supplementary** Fig. 2).
iii. The structure of an apo HTNV-L WT hexamer, determined at a resolution ranging from 3 to 9 Å (overall resolution 3.6 Å), reveals a peculiar configuration of polymerases into a trimer of dimers that had never been reported for any RNA-dependent RNA polymerase described so far (**Fig. 1d, Supplementary** Fig. 3). The central dimer acts as a central anchor on which two external dimers bind symmetrically. While the central dimer displays a unique configuration, the two external dimers globally adopt the same organization as the isolated dimer, the whole hexamer keeping the two-fold C2 symmetry.

### Domain organization of HTNV-L

The HTNV-L polypeptide chain begin with the N-terminal ENDO domain and includes residues 1 to 220 (**Fig. 2a and b**). Its folding is relatively conserved compared to the isolated ENDO X-ray crystal structure that was comprising the residues 1 to 179^16^, except for residues 1-19 that adopt a new conformation due to the presence of the residues 180 to 220 (rmsd 0.9 Å on 123 Cα residues, **Supplementary** Fig. 4).

**Figure 2.**
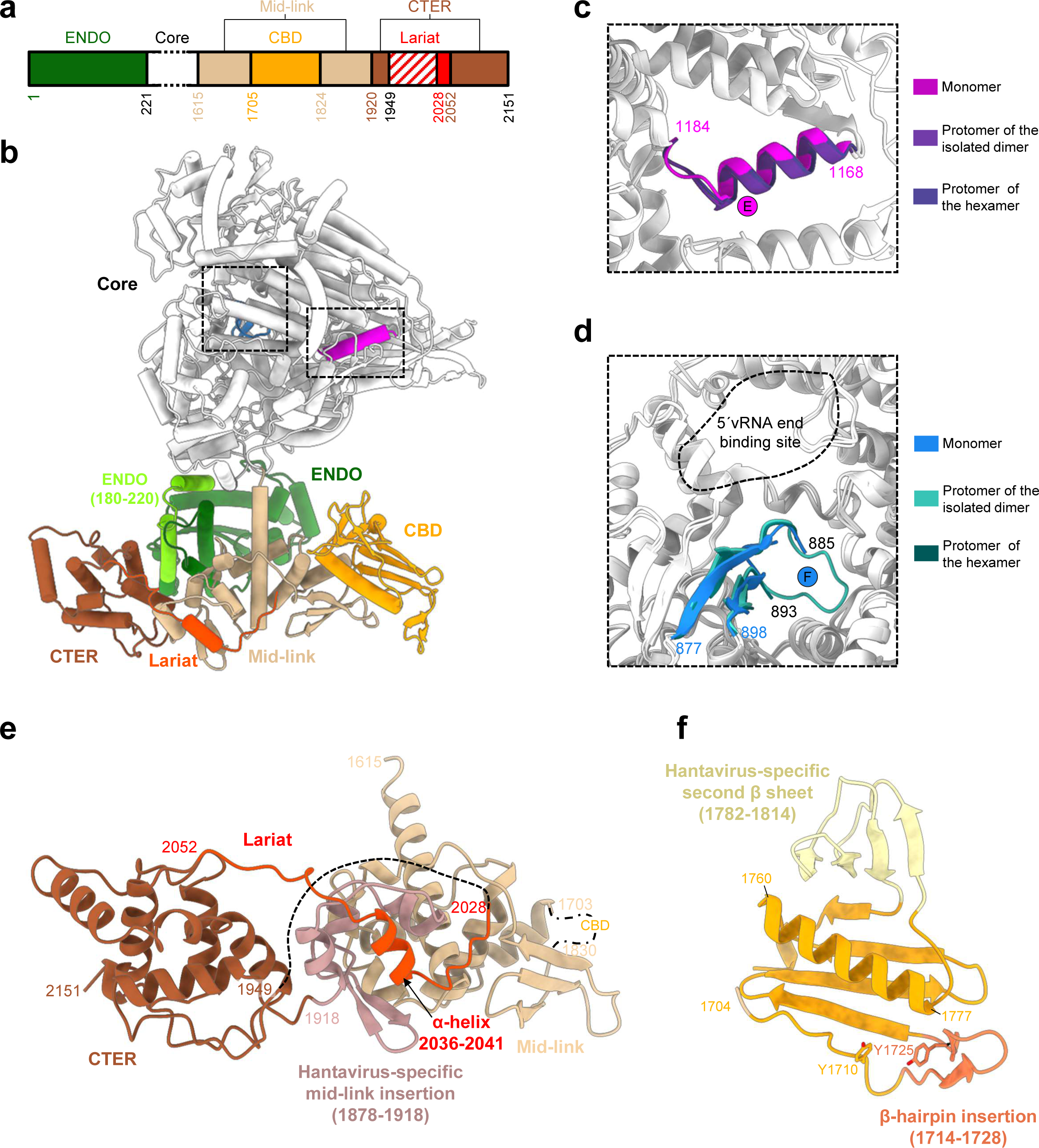
Protomer organization of HTNV-L. **a** Schematic representation of HTNV-L domain structure. **b** Cartoon representation of HTNV-L protomer with the ENDO, the core, the mid-link, the CBD, the lariat and the CTER respectively colored in forest green, white, beige, orange, red and brown. The motif E and F are respectively colored in magenta and blue. **c** Zoom on the motif E colored in magenta for apo HTNV-L monomer, in purple for HTNV-L dimer and in blue for HTNV-L hexamer. **d** Zoom on the motif F colored in different shades of blue for apo HTNV-L monomer, dimer and hexamer. The 5′vRNA end binding site is identified with a dotted line. **e** Zoom on the mid-link, the CTER and the lariat colored as in **a.** The Hantavirus-specific mid-link insertion is indicated and colored in light pink. The CBD position is indicated with a dashed dotted line. The dotted line that links the CTER to the lariat represents the residues of the lariat that are not visible due to flexibility. **f** Zoom on HTNV-L CBD model predicted by Alphafold and validated by a 8 Å cryo-EM map. The central 5-stranded β-sheet and the α-helix that are common to all sNSV CBD is shown in orange cartoon. The β-hairpin insertion that is likely to be essential for cap binding is shown in dark orange. The tyrosines that are predicted to bind the cap are shown as stick. The Hantavirus specific second β-sheet is shown in light yellow.

Following the ENDO is the polymerase core (221-1601), which adopts a conformation equivalent to the previously obtained apo HTNV-L_D97A_ structure^8^ (rmsd between HTNV-L_WT_ and HTNV-L_D97A_ monomer: 1.0 Å on 1275 Cα residues). The overall resolution improvement, from 3.1 Å HTNV-L_D97A_ to 2.6 Å in HTNV-L_WT_, results in a more accurate definition of several regions, notably a β-hairpin of the core-lobe region (residues 676-695) that, by analogy with other Bunyavirus polymerases, could be involved in the stabilization of the C-terminal region (**Supplementary** Fig. 5). The comparison of all HTNV-L apo protomer cores reveal their structural similarity, independently of their oligomeric states. Their catalytic motifs display an inactive configuration, with a motif E organized in an α-helix, as previously reported for monomeric apo HTNV-L_D97A_ structure^8^ **(Fig. 2c)**. One exception is the motif F tip, which is disordered in HTNV-L monomer and in the protomers of HTNV-L hexamer, but ordered in isolated HTNV-L dimer. Intriguingly, the position of the motif F in isolated HTNV-L dimer is closing the template entry tunnel, further preventing HTNV-L dimer activity (**Fig. 2d, Supplementary** Fig. 6). The putative priming loops also slightly rearrange themselves depending on the oligomeric states. These loops are named according to their role in replication initiation in Influenza polymerases^21^, but are located away from the active site in HTNV-L, where they rather correspond to template exit plugs^8^. Their exact position is conserved in HTNV-L monomers and HTNV-L hexamers, but differs in HTNV-L dimer. Their conformational change is associated with the reorganization of the thumb-ring residues 1455- 1463 from an α-helix in HTNV-L monomer/hexamer to a loop in HTNV-L isolated dimer (**Supplementary** Fig. 7).

After the polymerase core lies the C-terminal region that is divided in three structural domains: the mid-link (residues 1615-1704 and 1824-1919), the CBD (1705-1823) and the CTER (residues 1920-1949, 2053-2151) (**Fig. 2a**). From the CTER domain protrudes a long loop that will be called lariat region, in reference to the name given of a similar protrusion in DBV- L^22^ (residues 1950-2052, ordered part 2028 to 2052) (**Fig. 2a and e**).

The mid-link contains an α-helical mid region and a 3-stranded β-sheet link. While the mid region has a size comparable to equivalent region in other Bunyavirus polymerases^10,12,22,23^, the HTNV-L link encloses a hantavirus-specific insertion containing 3 additional small α-helices and a β-hairpin (**Fig. 2e**, **Supplementary** Fig. 8).

Inserted in the mid-link lies the CBD. As it does not interact with any other domain in HTNV-L dimers and hexamers, the CBD remains highly flexible precluding its structural characterization at high-resolution. Nonetheless, the 8 Å density enables an unambiguous fit of the CBD model that was predicted with high confidence by Alphafold (per-residue confidence score – pLDDT - comprised between 67 and 90, with an average score of 80, on a scale from 0 to 100) (**Fig. 2f, Supplementary fig. 9)**. The predicted CBD model consists in a central 5-stranded β-sheet containing a small β-hairpin insertion (1714-1728) between the first and the second strand. A long α-helix (residues 1760-1777) packs against the central β- sheet. In addition to this minimal fold that is found in all sNSV CBD structures determined so far ^10,12,22,24–26^, a second 4-stranded β-sheet is predicted that is specific to Hantaviruses (**Fig. 2f**, **Supplementary** Fig. 8). The capability of sNSV CBD to bind capped RNA has been related to the length of the β-hairpin insertion^26^. HTNV-L CBD predicted model displays a length comparable with the one of LACV-L CBD and DBV-L CBD^10,22,23^, suggesting that HTNV-L may be capable of cap binding in certain conditions. The resolution of the CBD structure is not sufficient to identify the residues involved in cap binding, however the comparison of HTNV CBD model with the structures of other sNSVs CBD suggests that the residues Y1710 and Y1725 may stack the cap m^7^GTP moiety (**Fig. 2f, Supplementary** Fig. 8). In the low-resolution CBD density of HTNV-L dimer, the β-hairpin insertion however appears to be disordered (**Supplementary** Fig. 9), suggesting that large conformational change of the C-terminal regions are likely to be necessary for β-hairpin stabilization and subsequent cap-binding, as observed for LACV-L^10,11^.

Finally, the CTER domain is composed of 6 α-helices that intertwine and from which protrudes a long and extended loop called the lariat (residues 1950-2052, ordered part 2028 to 2052) (**Fig. 2e)**. The lariat α-helix 2036 to 2041 interacts with the α-helical part of the hantavirus-specific insertion of the mid-link, strengthening the mid-link/CTER interaction.

### HTNV-L protomers can adopt two drastically different conformations

When analysing the protomers that are composing the different dimers, it is striking to observe that they adopt two very different configurations (**Supplementary Movie 1**). The protomers that are composing the isolated dimer and the external protomers of the hexamer globally adopt the same conformation, named conformation A, whereas the protomers of the central dimer of the hexamer adopt another configuration, named conformation B (**Fig. 1d and 3**). Superimposing the cores of conformers A and B reveals that the orientation of their respective ENDO differs by 68° (**Supplementary Movie 1**). Their entire C-terminal region also differs in position, with the mid-link domain that acts as a pivot point and modifies its orientation by 83° between conformers A and B (**Fig. 3a, Supplementary Movie 1**). Within the C-terminal region, the relative position of the domains also differs between the two protomer conformations: superimposing the mid-link of protomers A and B reveals that their CTER differs both in orientation (79°) and position (50 Å apart), while the CBD position differs by around 25 Å (**Fig. 3b**). As a result, the ENDO and the C-terminal region of protomers A and B exhibit very different mode of interaction with the core domain. The ENDO of both conformers interact with the lid domain, although this interaction involves different residues (**Supplementary** Fig. 10). The C-terminal region of conformer A is located away from its polymerase core, whereas the core and the C-terminal regions of conformer B tightly packs with its polymerase core, with the mid-link binding to the lid domain and the CTER interacting with the thumb and the thumb-ring domains (**Fig. 3b, Supplementary** Fig. 10).

**Figure 3.**
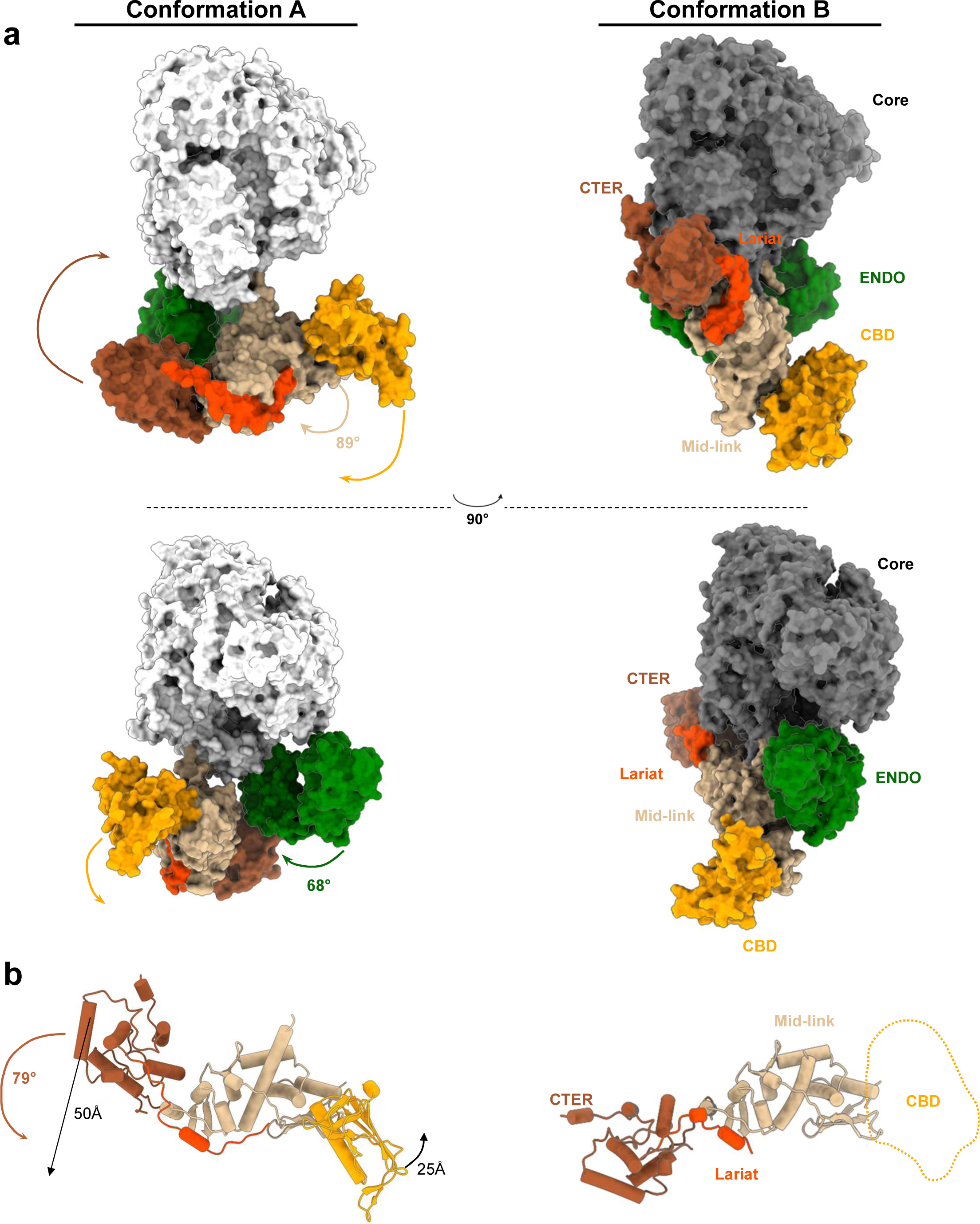
Comparison of the two different conformations of the protomers. **a** Surface structure colored as in Fig. 2 showing the conformation of protomer A, present in isolated dimer and in the external dimers of the hexamer, and protomer B, present in the central dimer of the hexamer. The rotations between both conformations are indicated. Two views that differ by 180° are displayed. **b** Zoom on the conformation of the C-terminal region of conformers A and B. Domain rotations and translations are indicated. A dotted line represents the position of the CBD of the central dimer.

### Presence of two different conformations of the protomers results in the formation of two different symmetric dimers

The divergence in the configurations of conformers A and B is compatible with the formation of two very different stable dimers.

The association of two protomers in conformation A results in the formation of dimer A that can be found either isolated or at the periphery of the hexamer (**Fig. 1c and d**). The solvent accessible surface of one protomer buried at the dimer’s interface, as estimated by PISA^27^, is about 4200 Å^2^, indicating a stable and intricate interaction involving numerous binding regions. The central hubs that promote dimer A formation are the CTER domains (**Fig. 4a and b**, **Table 1**). Of crucial importance is the extreme HTNV-L C-terminal loop containing the residues 2145 to 2151 that swaps into a hydrophobic groove of the opposite CTER (**Fig. 4b**), forming specific interactions that notably involve the two last HTNV-L residues F2150 and Y2151. F2150 is located in a hydrophobic cleft of the facing protomers where it stacks with P2120 and Y2064 (**Fig. 4b**). The aromatic ring of the C-terminal residue Y2151 forms hydrophobic interactions with the aliphatic moieties of N2110 and R2063, and the hydroxyl group of Y2151 makes hydrogen bonds with R2063 and D2106 (**Fig. 4b**). In addition, the interaction between CTER domains is reinforced by the binding of R2139 from protomer 1 with C2119 and D2121 of protomer 2 (**Fig. 4b and c**). Importantly, all the residues involved in the interactions between the CTER domains are conserved amongst Hantaviruses (**Supplementary Data 1**), suggesting that this interaction is likely to be biologically relevant and maintained during evolution.

**Figure 4.**
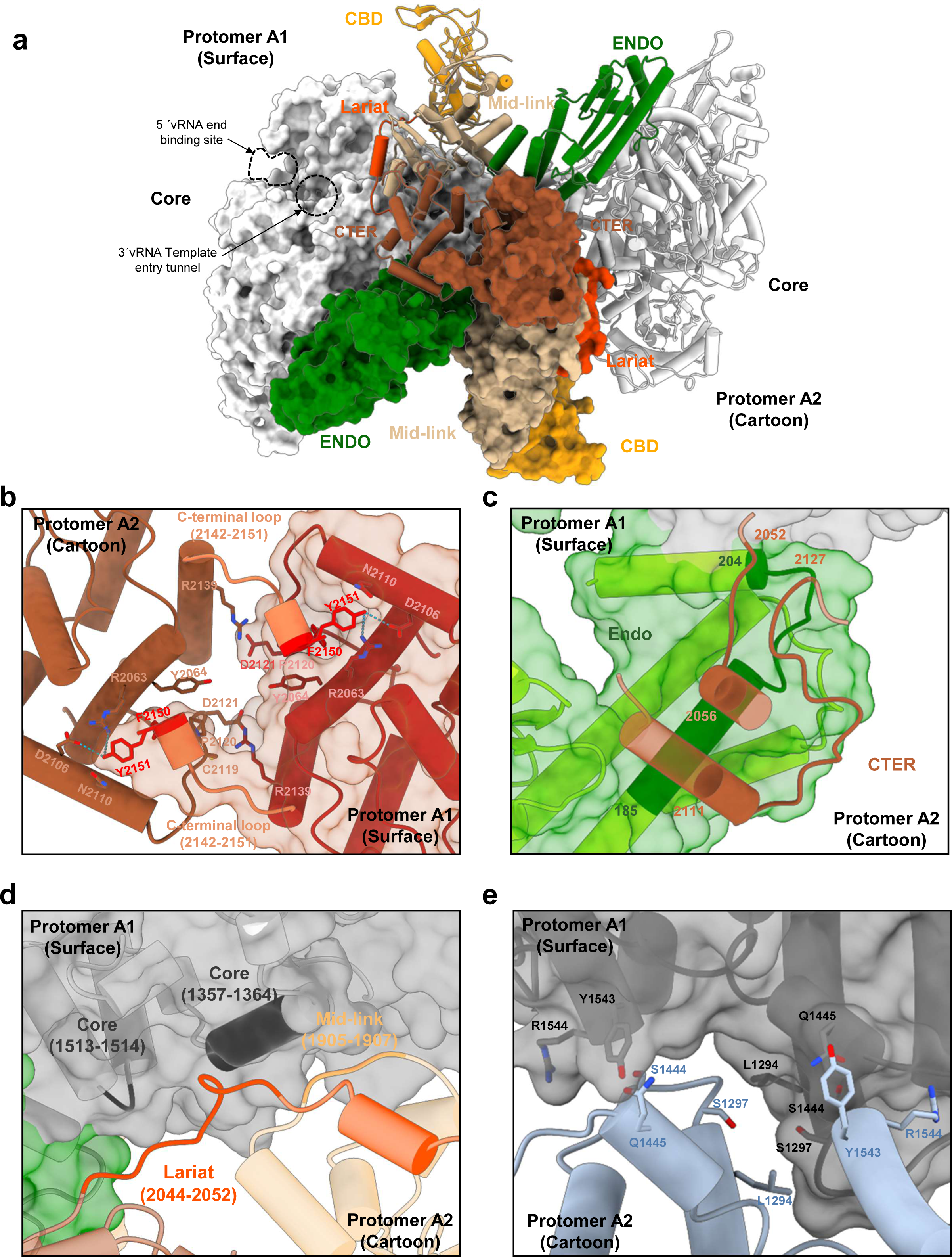
Dimer organization and protomer interactions. **a** Dimer structure with protomer A1 shown as surface and protomer A2 shown as cartoon. The domains are colored as in Fig. 2. The 5′vRNA end binding site and the template entry tunnel, to which the 3′vRNA bind, are indicated. **b** Zoom on the interaction of the CTER domains in the same orientation as in **a**. Protomer A1 surface is shown in transparent dark red and its cartoon representation in dark red. Protomer A2 is shown as brown cartoon. Residues in interaction are shown as stick. The C-terminal loops 2142-2151 that are crucial in the C-terminal swapping stabilization are shown in coral with F2150 and Y2151 shown in red. **c** Zoom on the interaction of the ENDO of protomer A1 with the CTER of protomer A2. Protomer A1 surface and cartoon are colored as in **a** with the regions that interact shown in dark green with residue numbers indicated. The CTER regions of protomer A2 that interact with protomer A1 are displayed as brown cartoons, the regions that interact are indicated. **d** Interaction of the lariat and the mid-link of protomer A2 with the core of protomer A1. Both protomers are colored as in **a** and shown as semi-transparent. The regions of the lariat, the mid-link and the core that interact are respectively shown in dark orange, orange and dark gray. Residue numbers of the regions that interact are indicated. **e** Interactions of the core of protomer A1, shown in grey cartoon and surface, with the core of protomer A2, shown as a light blue cartoon. The interacting residues are shown as sticks and are numbered.

**Table 1:**
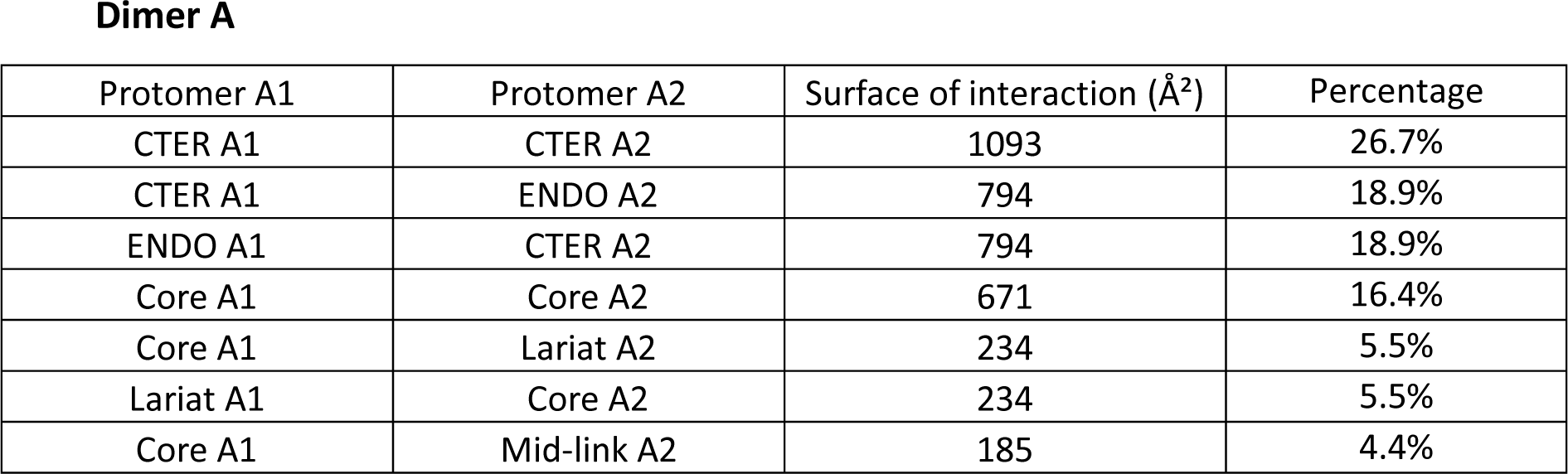
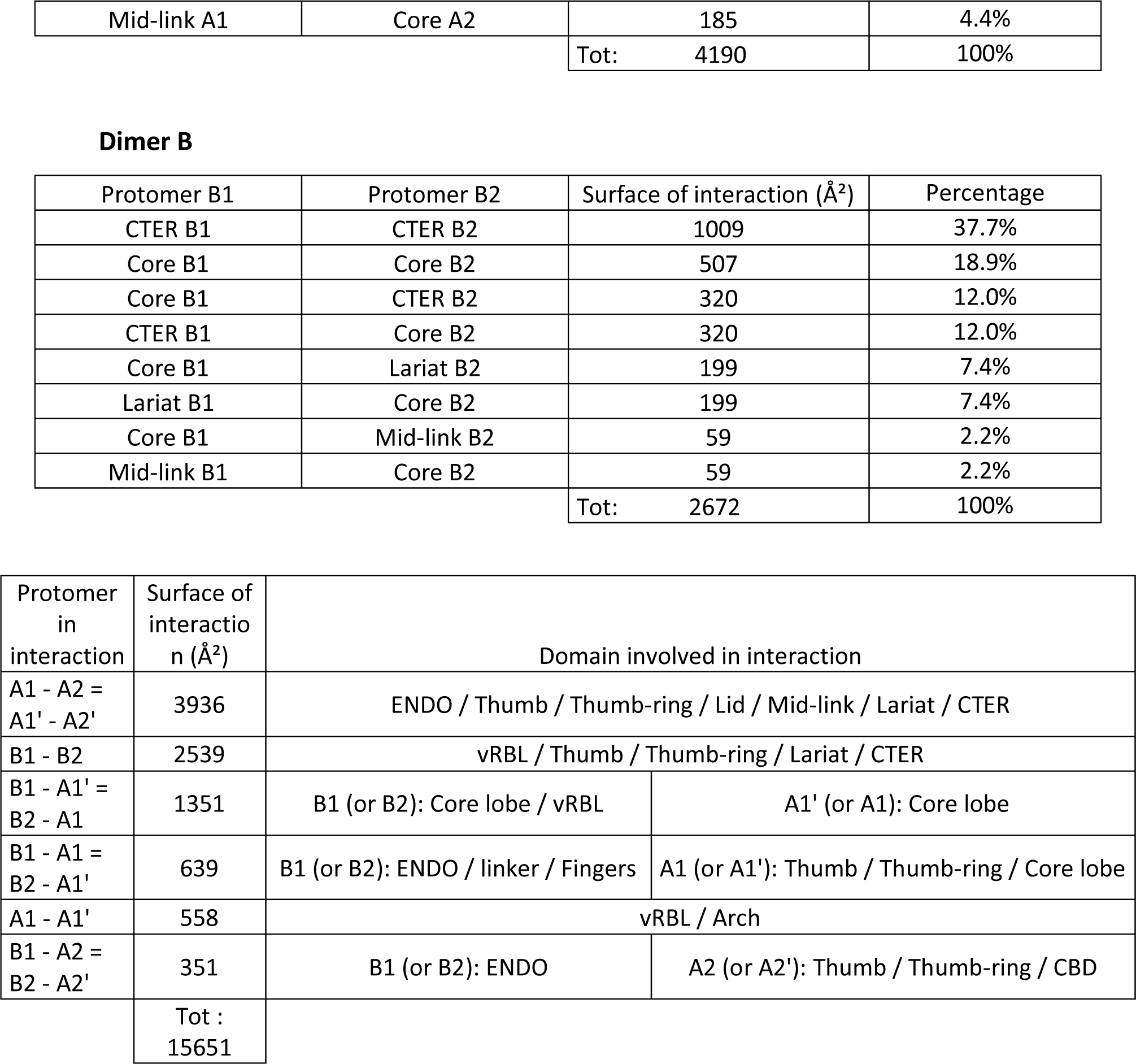
Analysis of the interaction surfaces within multimers. **a, b** The domains that interact within protomers of dimer A (**a**) and protomers of dimer B (**b**) are indicated along with their interacting surfaces. Percentages quantify each specific interaction compared to the global interaction between protomers. **c** List of the interactions between protomers in the context of the hexamer. Protomers are named according to **Fig.5b**. Their interaction surface is indicated, as well as the domains involved.

The CTER domains are also involved in stabilizing the different domains that compose the dimer A. The CTER residues 2052-2056 and 2111-2127 from one protomer are interacting with the ENDO residues 185-204 of the other protomer, leading to a mutual stabilization of these regions (**Fig. 4c**, **Table 1**). In addition, the lariat residues 2044-2052, together with the mid-link 1905-1907 residues, make hydrophobic interactions with the core residues 1357-1364 and 1513-1514 of the other protomer (**Fig. 4d**, **Table 1**).

The only interaction between the two HTNV-L monomers that do not involve the CTER are mediated by few residues of the core. A symmetric interaction is notably observed between L1294 from one monomer with S1297 from the other, while S1444 and Q1445 interact with Y1543 and R1544 (**Fig. 4e**, **Table 1**). The buried interface remains minimal (16%) and the residues that interact are not conserved amongst Hantaviruses, suggesting the limited impact of CORE-CORE interaction in dimer formation.

This intricate organization is stabilizing most of the domains, except few regions that are located outwards of the dimer, such as the CBD and the external part of the ENDO. These domains display a higher degree of mobility and were not clearly discernible in the global 3D reconstruction map. Hence, image processing involving signal subtraction, masking and local refinement were necessary to visualize their organization (**Supplementary** Fig. 2).

The symmetric dimer A can be compared with the symmetric dimer generated by the interaction of protomers in conformation B. This second type of stable dimer constitutes the central dimer of the hexamer (**Fig. 1d**). In dimer B, each protomer buries about 2700 Å^2^ of solvent accessible surface at the dimer’s interface, which is less than the one in dimer A, but still significant (**Table 1**, **Fig. 5a**). Despite the large differences in the arrangement of the protomers present in dimers A and B, it is interesting to note the presence of some similarities in the regions that contribute the most to the interface. In both cases, the CTER domains remain the main hub of the protomer interactions (**Fig. 5a**, **Table 1**), that notably involve the swapping of C-terminal residues 2142-2151 into the CTER hydrophobic groove of the second protomer (**Fig. 3b and 5a**). However, some protomer-protomer interactions are specific to the dimer B. For instance, the CTER α-helix residues 2131-2141 and the CTER loop 2118-2121 of one protomer B interacts with their symmetrical equivalent. Moreover, the CTER being in contact with the thumb-ring and the thumb is bolstered through interactions with the core of the second protomer B (**Fig. 5a**, **Table 1)**. In addition, the polymerase cores of the two protomers B make hydrophobic contacts through their respective residues 1301-1303 and 1252-1256, although their contribution to the dimer’s interface remain modest (19%). Finally, the lariat residues interact with the polymerase core of the other protomer and in particular its viral RNA binding-lobe (vRBL) (**Table 1**).

**Figure 5.**
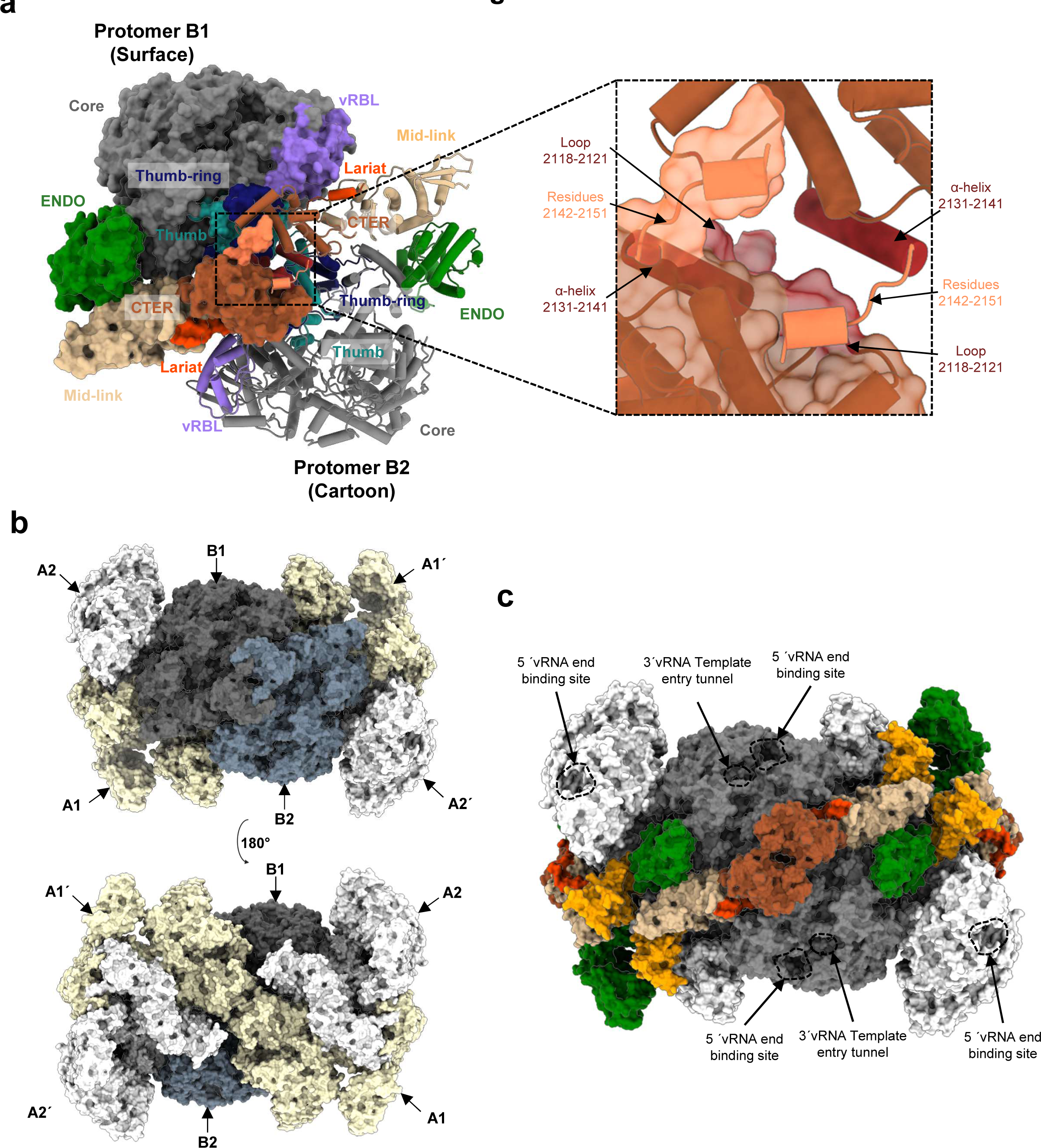
Hexamer structure: interactions between protomers. **a** Interaction of two protomers B results in the formation dimer B that corresponds to the central dimer of the hexamer. One protomer is shown as a surface and the second as a cartoon. The ENDO, the mid-link, the CTER and the lariat are colored as in Fig. 2. The vRBL, the thumb and the thumb-ring are respectively colored in purple, light sea green and midnight blue. On the right side, zoom on the CTER. Interacting residues of the CTER are indicated. Residues 2142-2151 that swap into the other protomer are displayed in orange. **b** Surface representation of the hexamer visualized in the same orientation as in **a** (top) and rotated by 180° (bottom). The protomers B1 and B2 are colored as in **a,** the external protomers A1 and A1’ are colored in white, the external protomers A2 and A2’ in beige. **c** Surface of the hexamer structure with domains colored as in Fig. 2. The cores of the protomers of the central dimer are colored in grey while the cores of the protomers of the external dimers are colored in white. The 5′vRNA end binding site and the template entry tunnel, to which the 3′vRNA bind, are indicated.

### Interactions between HTNV-L dimers A and B result in the formation of HTNV-L hexamer

Dimer B is observed only when capped by two dimers of type A (**Fig. 5b**). Binding of dimers A to dimer B is directional, the same side of both external dimers A being in contact with the central dimer B, and induces minimal reorganisation of dimers A in comparison with their conformation in the isolated dimer form (**Supplementary** Fig. 11).

The symmetric hexamer formed cannot be further elongated into larger complexes using the known protomer conformation and the same dimer A – dimer B interface. One can however imagine that some sub-complexes could exist, that would contain less than six subunits. This would be consistent with the observation of trimers and tetramers visualized by mass photometry (**Fig. 1a**).

For complete hexamers, the interaction between protomers is extensive, reaching about 15.600 Å^2^ (**Fig. 5b** and **Table 1**). Analysis of the interface area between protomers confirms that the hexamer is indeed composed of a trimer of dimers as dimer A and both dimers B display large protomer interfaces, respectively around 3900 Å^2^ and 2500 Å^2^, whereas the other protomer-protomer interfaces found in the hexamer are smaller, comprised between 350 to 1310 Å^2^ (**Table 1**). However, the interface between the two types of dimers reaches 2300 Å^2^, supporting the stability of the hexameric form.

### Binding of viral RNA induces conformational changes and disrupt HTNV-L oligomers

In the hexamer, the ENDO and the CTER domains are positioned towards the interior part of the complex and are in tight interaction, whereas the 5′ and 3′ vRNA binding sites are solvent exposed (**Fig. 5c**). In the dimer, the 5′ and 3′ vRNA binding sites are also accessible (**Fig. 4a**). To analyze if HTNV-L symmetric multimers could therefore bind vRNA, HTNV-L was incubated with either 5′vRNA, or a combination of both 5′ and 3′vRNA with mutations introduced on the 5′vRNA end, referred to as “5′mut”, to promote 5′ and 3′ concomitant binding to HTNV-L^8^. Intriguingly, cryo-EM 2D class averages revealed that only 0.5% of symmetric multimers remain when 5′vRNA is added, the extremely low number of particles preventing further structural characterization. In presence of 5′mut and 3′vRNA, only monomers were visualized. This strongly differs with the 2D class averages visualized in the absence of vRNA in which 12.3

% of the particles were multimeric (**Fig. 6a**). To visualize the potential conformational changes that could explain oligomer disruption, 5′vRNA-bound HTNV-L structures were determined at an overall resolution of 2.8 Å (**Supplementary** Fig. 12**, Supplementary Table 2)**. 3D classification revealed the presence of two distinct conformations that will be called “intermediate” and “active” (**Fig. 6b**). In the intermediate conformation, 5′vRNA-binding induces local reconfiguration, notably in the motif F and in the linker connecting both the core- lobe and the finger domains. In addition, whereas very few global movements of HTNV-L are visible (**Supplementary** Fig. 13), 5′vRNA-binding induces several movements of the canonical RdRp motifs. The motif E retains its amphipathic α-helix configuration but changes its orientation compared to its position in apo HTNV-L (**Fig. 6b**). This alteration modifies the van der Walls interactions between motif E and the α-helix 929-948, inducing a global movement of the α-helix 929-948 and the ordering of the putative PR-loop. Minor movements are also observed in motifs A, C and D, although no magnesium ion is visible in the active site.

**Figure 6.**
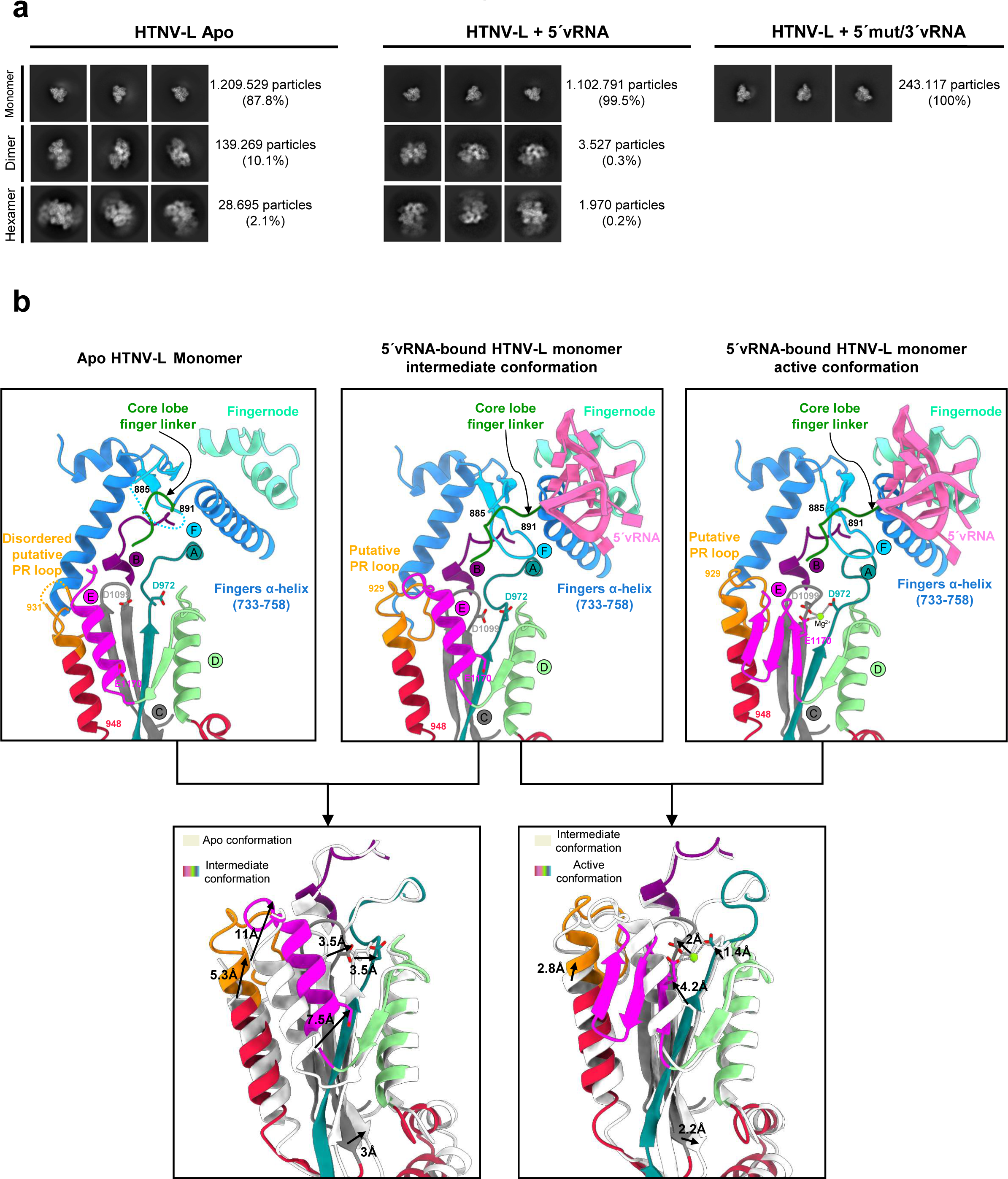
Disruption of HTNV-L oligomers induced by vRNA binding. **a** 2D class averages of HTNV-L apo (left), 5′vRNA-bound HTNV-L (middle) and 5′/3′RNA-bound HTNV-L (right). Monomer (top), dimer (middle) and hexamer (bottom) class averages are shown. The number of particles and the percentage compared to the entire dataset is indicated. **b** Zoom on the active sites of monomeric apo HTNV-L on the top left, 5′vRNA-bound intermediate conformation in the top middle and 5′vRNA-bound active conformation on the top right. The motifs, the putative prime-and-realign loop (PR loop), the 5′vRNA-binding site are labeled and have specific colors. Superimposition of monomeric apo HTNV-L with 5′vRNA- bound intermediate conformation (bottom left) and superimposition of 5′vRNA-bound intermediate conformation with 5′vRNA-bound active conformation (bottom right) are shown to highlight motif movements.

The reconfiguration of 5′vRNA-bound HTNV-L into a conformation compatible with activity induces a more drastic global opening of the polymerase which is associated with the large reorganization of the motif E from an amphipathic α-helix to a 3-stranded β-sheet (**Supplementary** Fig. 13). Whereas movements within motif E are significant, with large movements of each of its amino acid, the movements of the other motifs remain relatively small compared to the intermediate conformation (**Fig. 6b**). The reorganization of motif E induces the formation of a pocket and the binding of a magnesium ion that is coordinated between D972 of motif A, D1099 of motif C and E1170 of motif E (**Fig. 6b**).

The equilibrium between monomer, dimer and hexamer observed in the absence of vRNA is displaced by 5’vRNA binding towards the monomeric state (**Fig. 7**). The visualization of the equilibrium between different oligomers in the absence of RNA at the concentrations used in cryo-EM and mass photometry suggests that the equilibrium dissociation constant of the interaction is roughly in the micromolar range. Therefore, it is likely that small structural perturbations of the polymerase core could induce oligomer disruption. When superimposing the 5’vRNA-bound monomeric HTNV-L structure onto the HTNV-L isolated dimeric structure (**Supplementary** Fig. 14), we observe that the global opening of HTNV-L core induced by 5’vRNA-binding indeed modifies the orientation of the thumb, the thumb-ring, the lid and the bridge domains. As a result, clashes appear between the bridge domain of one protomer and the lariat region of the second, that are likely to be sufficient to displace the equilibrium towards the monomeric state. Altogether, these observations indicate that 5′vRNA binding results in the disruption of HTNV-L oligomers.

**Figure 7.**
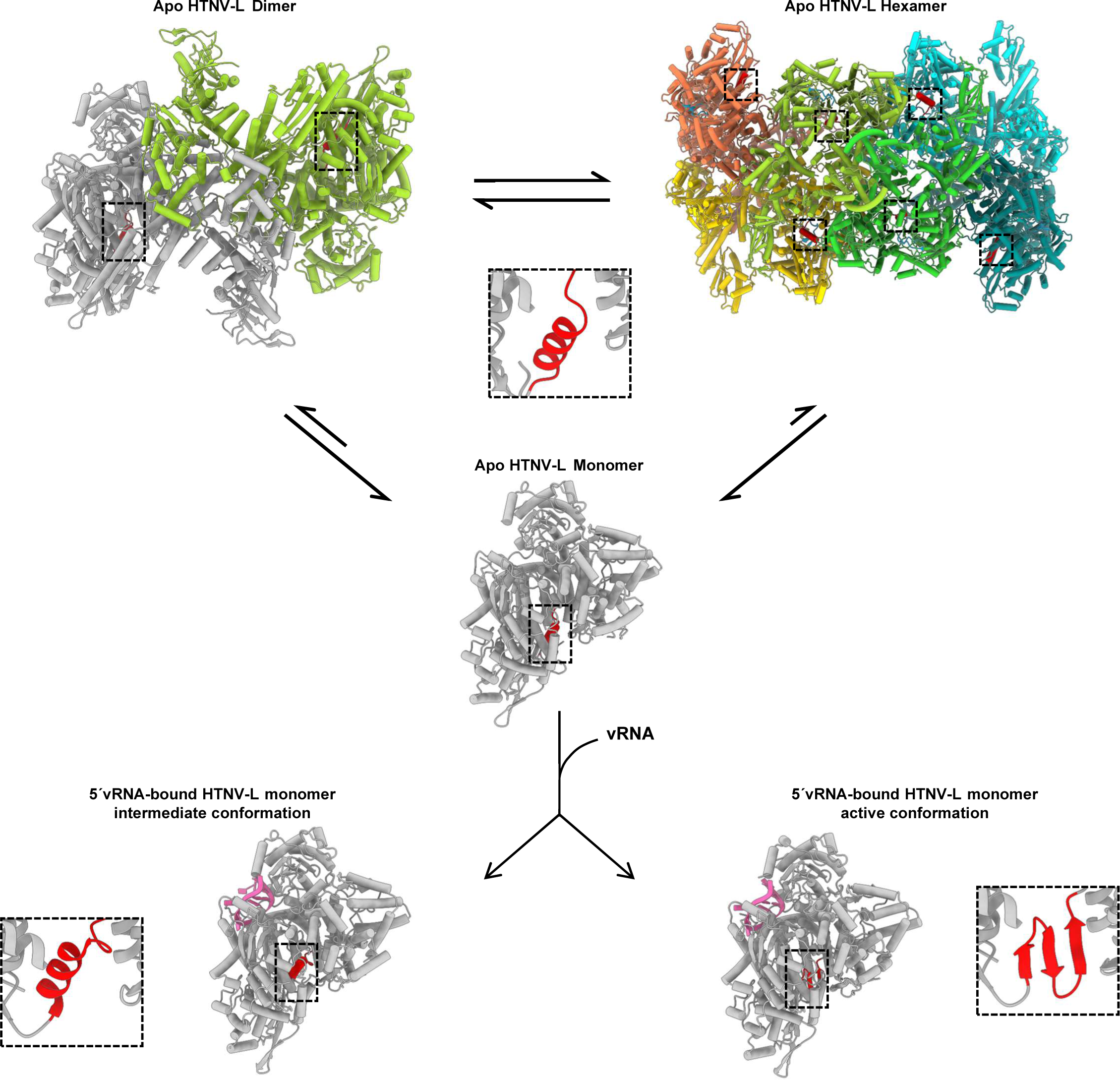
Disruption of apo HTNV-L multimers upon vRNA-binding is associated with HTNV-L conformational changes. HTNV-L apo is in equilibrium between dimers, hexamers and monomers as shown by cartoon representation of HTNV-L colored by protomers. Arrows indicate the equilibrium between the different oligomers. Motif E is shown in red and surrounded by a dotted line. In each protomer of HTNV-L apo, motif E is in an α-helical conformation as shown by a zoom where motif E is colored in red. vRNA binding disrupts the oligomers and results in two different conformations of monomeric HTNV-L that are displayed as a white cartoon with the 5′vRNA displayed in light pink. The conformation of motif E in both conformations is shown in red and is surrounded by a dotted line. A zoom on motif E shows its conformation in each map.

## DISCUSSION

### Comparison of HTNV-L C-terminal region with other *Bunyavirales* polymerases

The structures of HTNV-L dimer and hexamer reveal the organization of the C-terminal region of *Hantaviridae* polymerase (**Fig. 1 and 2**). It is striking to observe that, although the C-terminal region is quite divergent in term of amino acid sequence amongst Bunyavirus polymerases, the C-terminal domains organization of all *Bunyavirales* polymerases described so far is rather conserved (**Supplementary** Fig. 15). Bunyavirus polymerases all contain a mid-link domain, that is positioned centrally and links the polymerase core to the C-terminal region. The CBD is inserted in the mid-link and is also present in all *Bunyavirales* polymerases determined so far. Even the CTER domains that display the largest variability amongst bunyaviruses exhibit a common very important feature: a large protrusion that is named β-hairpin strut in LACV-L, and lariat in BDV-L and HTNV-L. In LACV-L and BDV-L this protrusion stabilizes the relative position of the C-terminal region and the core by interacting with a specific β-hairpin present in the core-lobe^10,22^ (**Supplementary** Fig. 5). In HTNV-L the core-lobe β-hairpin is conserved, and one can hypothesize that the region of the lariat that is not visible in the HTNV-L current structures might bind to the core-lobe β-hairpin and stabilize the entire HTNV-L structure (**Supplementary** Fig. 5). In HTNV-L, the visible part of the lariat in addition stabilizes the mid-link and therefore contributes to the steadying of the entire C-terminal region.

Altogether, these observations clearly indicate that *Bunyavirales* polymerases have conserved a common organization during evolution, even in their C-terminal region that diverges the most in terms of sequence.

### Comparison of *Bunyavirales* polymerase dimers: diversity in structure and in function

The HTNV-L structures reveal the formation of two very different dimers that display large interface areas. Interestingly, both dimers involve symmetric interactions of their CTER domains, with the swapping of the C-terminal residues of one protomer into the second. The conservation of this binding mode suggests the importance of the CTER domains in polymerase dimerization. Interestingly, the involvement of the CTER domains and the C- terminal α-helix swap had already been observed in LACV-L dimers **(Supplementary** Fig. 16a)^10^. The physiological relevance of LACV-L dimers however remained to be ascertained as, although they had been observed in the asymmetric unit of the C2 crystal form, the crystal packing may had induced significant conformational changes. The HTNV-L observations thus reinforce the hypothesis that the involvement of the CTER domain in symmetric dimer formation is physiologic and might be conserved amongst Bunyavirus polymerases.

In contrast, comparison of HTNV-L and MACV-L dimers^12,14,15^ clearly indicate the difference in dimer configuration, although protomer interactions involve in both cases the ENDO and the thumb domain of the polymerase core **(Supplementary** Fig. 16b). Interestingly, 3′vRNA- binding in MACV-L results in the stabilization of the dimers that become the majoritarian specie, whereas vRNA-binding in HTNV-L destabilizes the multimers. In our opinion, this suggests that different types of dimers or multimers are likely to exist during the replication and transcription cycles, that would be related to different biological roles.

### HTNV-L hexamers: a peculiarity never observed for polymerases

Formation of polymerase hexamers had never been reported for any sNSV polymerases. To our knowledge, the only other report describing high-resolution structures of large polymerase multimers concern single-stranded positive sense Picornaviruses, such as Poliovirus or Foot-and-Mouth-disease virus, that replicate at the cytosolic surface of cytoplasmic membranes where they form planar and tubular oligomeric arrays^27,28^. These arrays correlate with cooperative RNA-binding and RNA elongation^29^. These multimers are thus in strong contrast with the observed HTNV-L hexamers that are visualized only in the absence of RNA. In addition, HTNV-L hexamers cannot be further elongated since the relative orientation of dimer A C2 axis and dimer B C2 axis is not compatible with the binding of other dimers. It remains to be investigated if the observed hexameric polymerase is specific to Hantaviruses or if it also exists in other *Bunyavirales* in specific conditions.

### Hypothetical role of HTNV-L dimers and hexamers

*In vitro* dimer and hexamer assemblies occur in buffers that are typical to structural and functional assays, pH 8 in presence of 250 mM NaCl, that are not far from physiological conditions. The visualization of the multimers *in vitro* opens the question of their presence and their function during cell infection. The field of research opened by our results is large and fascinating as Bunyavirus polymerases fulfil multiple functions during cell infection and therefore, as described in the precedent paragraph, are likely to form several types of dimers and multimers that would have different specific roles during the viral cycle. It would be interesting to identify if the symmetric dimers and hexamers observed here can become majoritarian in certain conditions and which elements or conditions would stabilize them. One can think of specific compartmentalization that would locally increase the polymerase concentration and would thus favour oligomerization. Another possibility is that host-cell co- factors may be necessary to stabilize these multimers. If symmetric dimers and hexamers observed here become majoritarian at one specific state, one possible role that we speculate here is a storage and protection state of newly translated polymerases that have not yet been in contact with viral RNA. Dimer and hexamer formation would prevent the large movements of the ENDO and the C-terminal region observed in HTNV-L monomers, thereby stabilizing the entire polymerase structure. The accessibility of the 5′ and 3′ vRNA binding sites to each protomer, and the observed disruption of multimers upon vRNA binding is compatible with this “polymerase storage” hypothesis.

If future research confirms the presence of these multimers during cell infection, our results would also provide a solid framework for future development of antivirals. Indeed, molecules that would stabilize and enrich the inactive multimer population could be considered as possible new leads that would preclude HTNV-L activation that is necessary for genome replication and transcription.

## MATERIAL AND METHODS

### Cloning, expression and purification

The full-length HTNV-L gene (strain 76-118/Korean haemorrhagic fever, GenBank: X55901.1 UniProt: P23456) flanked in 5′ by a sequence coding an N-terminal hexa-histidine tag was cloned in a pFastBac vector between NdeI and NotI restriction sites. The expressing baculovirus was prepared using the Bac-to-Bac method (Invitrogen)^31^. *Trichoplusia ni* High 5 cells were infected at 0.7x10^6^ cells/mL with 0.1 % v/v of baculovirus and harvested 120 h after infection. Culture medium was centrifuged at 1 000 g for 15 min. The cell pellets were resuspended in lysis buffer (30 mM HEPES pH8, 300 mM NaCl, 10 mM Imidazole, 2 mM TCEP and 5 % glycerol) supplemented with cOmplete EDTA-free protease inhibitor complex (Roche) and ribonuclease A (Roche). The lysate was sonicated during 3 min 30 s (10 s ON, 20 s OFF and 40 % intensity) and centrifuged during 45 min at 20 000 g and 4 °C. The supernatant was filtered at 0.8 µm and used for purification by nickel ion affinity chromatography (GE Healthcare). A washing step in the lysis buffer supplemented with 30 mM imidazole was followed by the elution in lysis buffer supplemented with 500 mM imidazole. HTNV-L fractions were slowly diluted by two at 4°C using the heparin buffer (30 mM HEPES pH8, 300 mM NaCl and 2 mM TCEP), loaded on a 1 ml heparin column (GE Healthcare) and eluted in the heparin buffer supplemented with 500 mM NaCl. For cryo-EM assays and some mass photometry assays, a final gel filtration step was performed in GF buffer (30 mM HEPES pH 8, 250 mM NaCl, 10 mM TCEP) using a S200 size exclusion chromatography column (GE Healthcare).

### Mass photometry

Mass photometry measurements were carried out on a OneMP mass photometer (Refeyn Ltd). Prior to sample preparation, the coverslip (No. 1.5H, 24x50mm, VWR) was washed several times with water and ethanol and dried with compressed air before being used as a support for the silicone gaskets (CultureWell^TM^ Reusable Gaskets, Grace Bio-labs). For mass/contrast calibration, 19 μL of GF buffer was deposited in a well of the silicone gasket and used to determine the focus. 1 μL of native marker (Native Marker unstained protein standard, LC0725, Life Technologies) diluted 400 x in the GF buffer was used for mass/contrast calibration that was monitored during 60 s using the AcquireMP software (Refeyn Ltd, Version 2.3.0) with a medium field of view (detection area of 56 µm^2^). For HTNV-L WT measurements, 19.5 μL of GF buffer was deposited on a well of the silicone gasket and used to find the focus.

0.5 μL of HTNV-L WT was added and gently mixed to reach a final concentration of 30 nM. Data acquisition was carried out as for mass/contrast calibration. All measurements were performed in triplicate. Data analysis was performed using DiscoverMP software (Refeyn Ltd, version 2.3.0).

### Electron microscopy

To obtain the structures of HTNV-L apo dimer and hexamer, 3.5 μL of HTNV-L at 1 µM was deposited on glow-discharged (25 mA, 45 s) UltraAuFoil 300 mesh R1.2/1.3 grids (Quantifoil). The grid was blotted for 3s (blot force 1) at 100 % humidity and 4°C in a Vitrobot Mark IV (Thermo Fisher Scientific) before plunge-freezing in liquid ethane.

To obtain the structure of 5′vRNA-bound HNTV-L monomers, HTNV-L at 1 µM was incubated with 10 µM 5′ vRNA (5′- UAG UAG UAG ACA CCG CAA GAU GUU A-3′) for 30 min at 4°C. Grid preparation was identical to the one described for HTNV-L apo.

To obtain 2D class averages of monomeric HNTV-L bound to 5′mut (5′- UAG GAG UAU CCA CCG CAA GA) and 3′ vRNA (5′ -UUU UGC GGA GUC UAC UAC UA-3′), HTNV-L at 1 µM was firstly incubated with 10 µM of 5′mut for 30 min at 4°C, and secondly incubated with 10 µM of 3′vRNA_1-25_ for 30 min at 4°C. Grid preparation was identical to the one described for HTNV-L apo.

To obtain 2D class averages of HNTV-L bound to 5′mut and 3′ vRNA, 2003 micrographs were collected on a 200kV Glacios cryo-TEM microscope (Thermo Fisher Scientific) equipped with a K2 summit direct electron detector (Gatan). Coma and astigmatism correction were performed on a carbon grid. Automated multi-holes (3x3) data collection was performed with SerialEM^32^. Movies containing 50 frames were acquired with a defocus ranging from -0.8 μm to -2.2 μm at a nominal magnification of 36.000x with a pixel size of 1.145 Å. The total exposure dose was 50 e^-^/Å^2^.

To obtain 3D structures of apo HTNV-L and 5′vRNA-bound HTNV-L, 14.650 and 26.745 micrographs were respectively collected on a 300kV Titan Krios cryo-TEM microscope (Thermo Fisher Scientific) equipped with a K3 direct electron detector (Gatan) and a Gatan Quantum LS energy filter^33^. Coma and astigmatism correction were performed on a carbon grid. Automated multi-holes (4x4) data collection was performed with EPU. Movies containing 40 frames were acquired with a defocus ranging from -0.8 μm to -2.0 μm at a nominal magnification of 105.000x with a pixel size of 0.839 Å. The total exposure dose was 40 e^-^/Å^2^.

### Image processing

To determine the structures of HTNV-L apo monomers/dimers/hexamers and HTNV-L 5′vRNA- bound monomers, micrograph movies were realigned using Motioncor2^34^ (**Supplementary** Fig. 2 and 3). The dose-weighted micrographs were imported into cryoSPARC 4.2.1^19^ for CTF determination using the “Patch CTF estimation (multi)” tool. High quality micrographs were selected with the “Manually Curate Exposure” tool and used for automated picking using the “Blob picker tool” utility. Different diameters of particles were used depending on the oligomeric state: between 90 and 190 Å for monomers, between 100 and 220 Å for dimers and between 100 and 300 Å for hexamers.

For the determination of apo and 5′vRNA-bound HTNV-L monomeric structures, a second 2D classification was performed, followed by an ab-initio reconstitution and a non-uniform refinement. For 5’vRNA-bound HTNV-L monomers, 3D classification without alignment with a global mask was used to separate the intermediate and the active conformations. Particles corresponding to each conformation were merged and used for a local refinement using a mask corresponding to HTNV-L core.

For the determination of HTNV-L structures in dimeric and hexameric states, the 2D class averages obtained with the blob picker or their corresponding particles were used for template and Topaz picking^20^. All the particles picked with blob picker, Topaz and template-picker that gave nice 2D class averages were used, following the removal of duplicates, to do an ab initio 3D reconstruction followed by a non-uniform refinement using the C2 symmetry. This provided global 3D reconstructions of the dimer and the hexamer.

To improve the density of the ENDO and the CBD of the dimer, symmetry expansion was done, followed by subtraction of the signal of one protomer. 3D classification without alignment focused either on the ENDO or the CBD provided one class in each case for which the ENDO or the CBD were more defined. A masked local refinement was then performed in each case.

Concerning the hexamer, to improve the density of the external dimer, a symmetry expansion was done followed by signal subtraction of a central protomer and an external dimer. A 3D classification without alignment separated particles with a better density of the external dimer that were used for local refinement.

Local resolution was estimated for the maps of HTNV-L apo monomers/dimers/hexamers and HTNV-L 5′vRNA-bound monomers and used to filter them.

To determine the 2D class averages of HNTV-L bound to 5′mut and 3′vRNA_1-20_, micrograph movies were imported in Relion 4.0.1^35^ in nine optic groups based the nine different beam shifts required for their acquisition. They were realigned using Motioncor2^34^ by applying the gain reference and the camera defect corrections. The micrographs were imported into cryoSPARC 4.2.1^19^ for the following steps. CTF parameters were determined using the “Patch CTF estimation (multi)” tool on the non-dose-weighted micrographs. Low quality micrographs were manually removed thanks to the “Manually Curate Exposure” tool. The selected micrographs were subjected to an automated picking with the “Blob picker tool” designed to select particles with a diameter comprised between 100 and 300 Å. The selected particles were extracted in a box size of 300x300 pixels^2^ and binned twice. Two rounds of 2D classifications were applied to remove contaminants and low-quality particles. At this stage only monomeric HNTV-L 2D classes were visible (**Supplementary** Fig. 1). Template picking and Topaz picking were performed using imported apo HTNV-L dimers and hexamers, but these procedures retrieved monomeric HTNV-L only.

### Model building in the cryo-EM maps

For the refinement of HTNV-L apo and 5’vRNA-bound monomers, HTNV-L apo core (PDB: 8C4S) and HTNV-L 5’vRNA-bound cores (PDB: 8C4T) were rigidly fitted in Chimera^36^. In Coot 0.9.8.1^37^, the restraint module was used to generate restraints at 4.3 Å that were used for flexible refinement to fit the main chain into density. The missing regions were manually built into COOT. Careful validation of every side chain position was performed in COOT before refinement in real-space using Phenix^38^.

For the refinement of HTNV-L dimer, HTNV-L ENDO crystal structure (PDB: 5IZE), HTNV-L WT apo core (described in this article) and an AlphaFold model^39^ of the mid-link, the CBD and the CTER domains of HTNV-L were rigidly fitted in Chimera^36^. The missing regions, in particular the lariat domain, the C-terminal region of the ENDO and missing loops of the core region were manually built into COOT. Careful validation of every side chain position was performed in COOT for every part except the CBD before being refined in real-space using Phenix^38^.

For the refinement of HTNV-L hexamer, the HTNV-L dimer structure was split in different rigid parts, namely the ENDO, the core, the mid-link associated with the lariat α-helix, the CBD and the CTER. These parts were used for rigid body fitting in the central dimer and in the most resolved external protomers. Careful validation of every side chain position was performed in COOT before being refined using Phenix real-space refinement^38^. For the less resolved external protomers, the map resolution was not sufficient for model building and the protomers were thus separated into isolated core, ENDO and C-terminal regions that were rigidly fitted in the focused map containing a central protomer and the external dimer. Interfaces between the less resolved external protomers and their interacting protomers were optimized in coot.

For each model, atomic model validation was performed with Molprobity^40^ and the PDB validation server. Model resolution according to the cryo-EM maps was estimated with Phenix at the 0.5 FSC cutoff. Protein-protein interactions were analysed with PISA^27^. Figures were created using ChimeraX^41^.

## DATA AVAILABILITY

The coordinates and structure factors generated in this study have been deposited in the Protein Data Bank and the Electron Microscopy Data Bank database under accession codes:

PDB ID 8QE5 EMD-18343 HTNV-L apo monomer

PDB ID 8QGT EMD-18390 5′vRNA-bound HTNV-L monomer in the intermediate state PDB ID 8QH3 EMD-18397 5′vRNA-bound HTNV-L monomer in the activated state

PDB ID 8QGU EMD-18391 chimeric map of HTNV-L dimer associated with the complete model of HTNV-L dimer

EMD-18392 cryo-EM map of HTNV-L dimer

EMD-18393 cryo-EM map focused on the ENDO of a protomer from HTNV-L dimer EMD-18394 cryo-EM map focused on the CBD of a protomer from HTNV-L dimer

PDB ID 8QHD EMD-18408 chimeric map of HTNV-L hexamer associated with the complete model of HTNV-L hexamer

EMD-18406 cryo-EM map of HTNV-L hexamer

EMD-18405 cryo-EM map of HTNV-L hexamer focused on one central protomer and two external protomers.

## ACKNOWLEDGEMENTS

We thank Guy Schoehn for setting up and maintaining the IBS/ISBG EM platform and for discussion; Eleftherios Zarkadas for support and technical advices on cryo-EM data collection on the IBS/ISBG Thermofisher Glacios; Lindsay Mcgregor and Romain Linares for data collection on the ERSF CM01 Krios; Martin Pelosse for technical advices on expression; Caroline Mas for training on the mass photometer; Aymeric Peuch for setting up and maintaining the EM computing cluster; Ambroise Desfosses for advices on cryo-EM image processing ; Daphna Fenel and Madalen Le Gorrec for technical support.

This work used the platforms of the Grenoble Instruct-ERIC center (ISBG; UAR 3518 CNRS-CEA-UGA-EMBL) within the Grenoble Partnership for Structural Biology (PSB) supported by FRISBI (ANR-10-INBS-05-02) and GRAL, financed within the University Grenoble Alpes graduate school (Ecoles Universitaires de Recherche) CBH-EUR-GS (ANR-17-EURE-0003). The electron microscope facility is supported by the Auvergne-Rhône-Alpes Region, the Fondation pour la Recherche Médicale (FRM), the fonds FEDER and the GIS-Infrastructures en Biologie Santé et Agronomie (IBiSA). We thank the platform staff who enabled us to perform these analyses. IBS acknowledges integration into the Interdisciplinary Research Institute of Grenoble (IRIG, CEA).

This work was supported by the ANR-19-CE11-0024 and the Institut Universitaire de France endowment to H.M.

## AUTHOR CONTRIBUTIONS STATEMENT

Q.D.T. and B.A. expressed and purified HTNV-L. Q.D.T. performed mass photometry measurements. H.M. and Q.D.T. collected cryo-EM data on a Thermo Fischer Scientific Glacios EM. Q.D.T. and B.A. performed cryo-EM image processing. B.A. determined the first cryo-EM reconstruction of HTNV-L dimers. Q.D.T. determined all the deposited structures. H.M., B.A. and Q.D.T. built the models based on the cryo-EM maps. H.M., Q.D.T., D.H. and B.A. performed structural analysis. H.M. supervises Q.D.T. and B.A. The project was conceived and used funding obtained by H.M. The manuscript was written by H.M., Q.D.T. and D.H. with inputs from B.A.

## COMPETING INTERESTS STATEMENT

The authors declare no competing interests.

## MATERIALS AND CORRESPONDANCE

Correspondence and material requests should be addressed to H.M.

## Notes

### Competing Interest Statement

The authors have declared no competing interest.

