## Supplementary Tables 1 and 2, Supplementary Figures 1 to 16 for "Structural characterization of the oligomerization of full-length Hantaan virus polymerase into symmetric dimers and hexamers"

Quentin Durieux Trouilleton et *al.*

#### Supplementary Table 1. Cryo-EM data collection, refinement and validation statistics of apo HTNV-L structures

| | HTNV-L apo monomer | HTNV-L apo dimer | | | HTNV-L apo hexamer | |
|---|---|---|---|---|---|---|
| | PDB 8QE5<br>EMD-18343 | PDB 8QGU<br>EMD-18391 (composite map) | | | PDB 8QHD<br>EMD-18408<br>(composite map) | |
| | | EMD-18392 (overall dimer) | EMD-18393 (refine ENDO) | EMD-18394 (refine CBD) | EMD-18406 (overall hexamer) | EMD-18405 (one central and two external protomers) |
| **Data collection and processing** | | | | | | |
| Microscope | Thermo Fisher Scientific Krios | | | | | |
| Camera | Gatan K3 | | | | | |
| Magnification | 105 000 | | | | | |
| Voltage (kV) | 300 | | | | | |
| Electron exposure (e-/A2) | 40 | | | | | |
| Defocus range (µm) | -0.8 to -2.0 | | | | | |
| Pixel size (Å) | 0.839 | | | | | |
| Initial/Final micrographs (no.) | 14650/11502 | | | | | |
| Symmetry imposed | C1 | C2 | C1 | C1 | C2 | C1 |
| Final particles (no.) | 142.391 | 139.269 | 56.805 | 53.788 | | |
| Map resolution (Å) | 2.6 | 3.0 | 3.1 (around 4 Å for the ENDO | 3.2 (around 8 Å for the CBD) | 3.2 | 3.6 |
| FSC threshold | 0.143 | 0.143 | | | 0.143 | |
| Map resolution range (Å) | 2.25-4.0 | 2.5-7.0 | 2.5-6.0 | 2.5-7.0 | 2.5-10.0 | 2.5-10.0 |
| **Refinement** | | | | | | |
| Model resolution (Å) 0.5 FSC threshold | 2.6 | 3.2 | | | | 3.3 |
| Map sharpening B factor (Å$^2$) | -80 | -80 | | | -80 | -80 |
| Model composition | | | | | Two internal protomers | One internal protomer and one external protomer |
| Protein residues | 1340 | 4064 | | | 3746 | 3872 |
| B-factor (Å$^2$, min-max (mean)) | 0.00-61.72 (19.56) | 23.64-257.21-(76.92) | | | 56.56-486.5-(157.97) | 7.76-246.25 (94.94) |
| R.m.s deviations | | | | | | |
| Bond lengths (Å) | 0.003 | 0.002 | | | 0.002 | 0.004 |
| Bond angles (°) | 0.513 | 0.503 | | | 0.517 | 0.6 |
| **Validation** | | | | | | |

| | | | | |
|---|---|---|---|---|
| MolProbity score | 1.95 | 1.63 | 1.81 | 1.89 |
| Clashscore | 7.16 | 7.82 | 9.99 | 10.79 |
| Poor rotamers (%) | 3.02 | 0.00 | 0.42 | 0.41 |
| **Ramachandran plot** | | | | |
| Favored (%) | 96.83 | 96.73 | 95.79 | 95.17 |
| Allowed (%) | 3.17 | 3.27 | 4.05 | 4.80 |
| Disallowed (%) | 0 | 0 | 0.16 | 0.03 |

#### Supplementary Table 2. Cryo-EM data collection, refinement and validation statistics of 5'-bound HTNV-L structures

| | **HTNV-L 5'-bound "intermediate"** | **HTNV-L 5'-bound "activated"** |
|---|---|---|
| | PDB 8QGT<br>EMD-18390 | PDB 8QH3<br>EMD-18397 |
| **Data collection and processing** | | |
| Microscope | Thermo Fisher Scientific Krios | |
| Camera | Gatan K3 | |
| Magnification | 105 000 | |
| Voltage (kV) | 300 | |
| Electron exposure (e-/A2) | 40 | |
| Defocus range (μm) | -0.8 to -2.0 | |
| Pixel size (Å) | 0.839 | |
| Initial/Final micrographs (no.) | 26745/18002 | |
| Symmetry imposed | C1 | |
| Final particles (no.) | 882.132 | 111.684 |
| Map resolution (Å) | 2.8 | 2.8 |
| FSC threshold | 0.143 | 0.143 |
| Map resolution range (Å) | 2.50-4.25 | 2.5-4.25 |
| **Refinement** | | |
| Model resolution (Å) 0.5 FSC threshold | 2.8 | 2.8 |
| Map sharpening B factor (Å$^2$) | -80 | -80 |
| Model composition | | |
| Protein residues | 1289 | 1309 |
| Nucleotide residues | 12 | 11 |
| Ligands | 0 | 1 |
| Water | 0 | 0 |
| B-factor (Å$^2$, min-max (mean)) | | |
| Protein | 0/64.18 (18.12) | 0/37.47 (8.45) |
| Nucleotides | 0/60.81 (12.44) | 0/37.83 (10.13) |
| Ligands | 0 | 10.34/10.34 (10.34) |
| R.m.s deviations | | |
| Bond lengths (Å) | 0.003 | 0.003 |
| Bond angles (°) | 0.502 | 0.506 |
| **Validation** | | |
| MolProbity score | 1.78 | 1.73 |
| Clashscore | 7.14 | 7.49 |
| Poor rotamers (%) | 2.11 | 1.55 |
| Ramachandran plot | | |
| Favored (%) | 97.24 | 96.99 |
| Allowed (%) | 2.76 | 2.93 |
| Disallowed (%) | 0 | 0 |

### Supplementary Figure 1

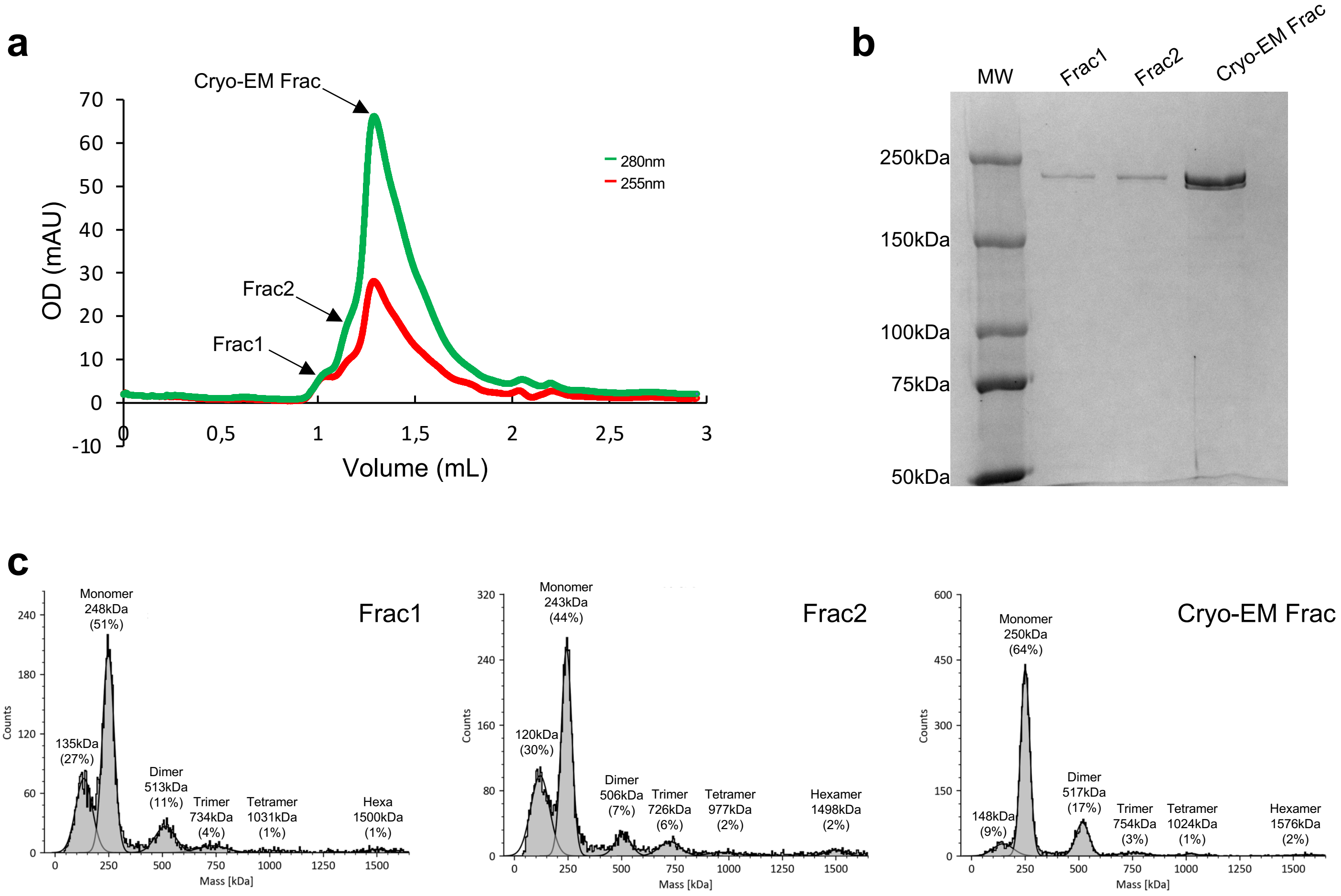


**Supplementary Figure 1 HTNV-L purification and characterization of its oligomerization**

**a** Size-exclusion chromatography profile on a superdex 200 increase 3.2/300 column. The optical density (OD) for 255 nm and 280 nm is indicated respectively in red and green as a function of the elution volume. The three fractions analyzed in **b** and **c** are indicated with arrows.

**b** SDS-PAGE gels of the 3 fractions indicated in **a**. The molecular weight marker (MW) is shown on the left.

**c** Mass photometry of the 3 fractions indicated in **a**. The mass in kDa and the percentage of each specie detected is indicated.

### Supplementary Figure 2

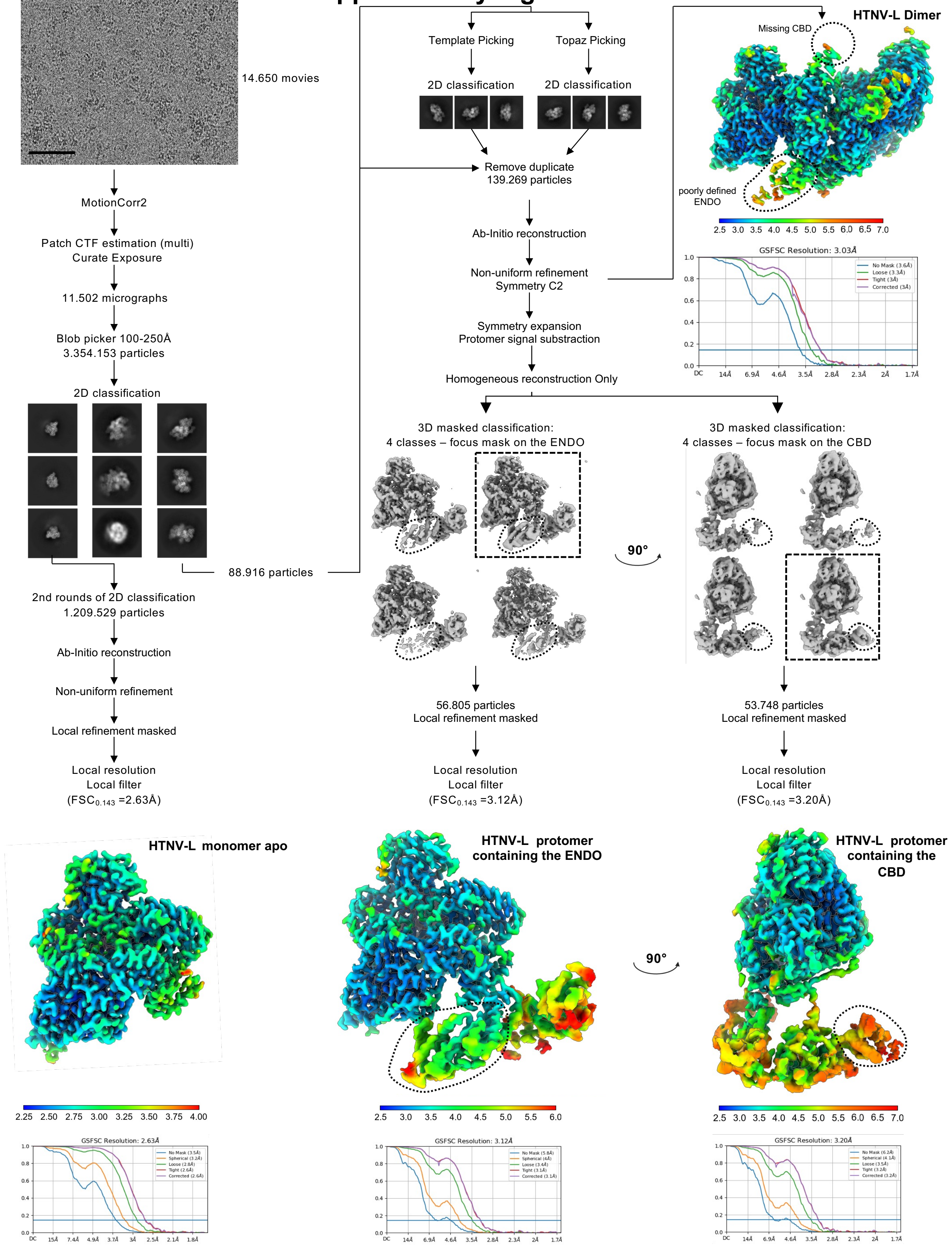


**Supplementary Figure 2 Image processing strategy to obtain monomeric and dimeric apo and HTNV-L**

A representative image of HTNV-L apo is displayed. The scale bar corresponds to 50 nm. The image processing workflow including 2D class averages, 3D class averages and the final reconstructions are displayed. Regions used for masking are indicated with a dotted line. 3D class averages chosen for further processing are surrounded by a rectangle dotted line. Electron density maps are colored according to the local resolution. Gold-standard Fourier Shell Correlation curves (FSC) are displayed.

### Supplementary Figure 3

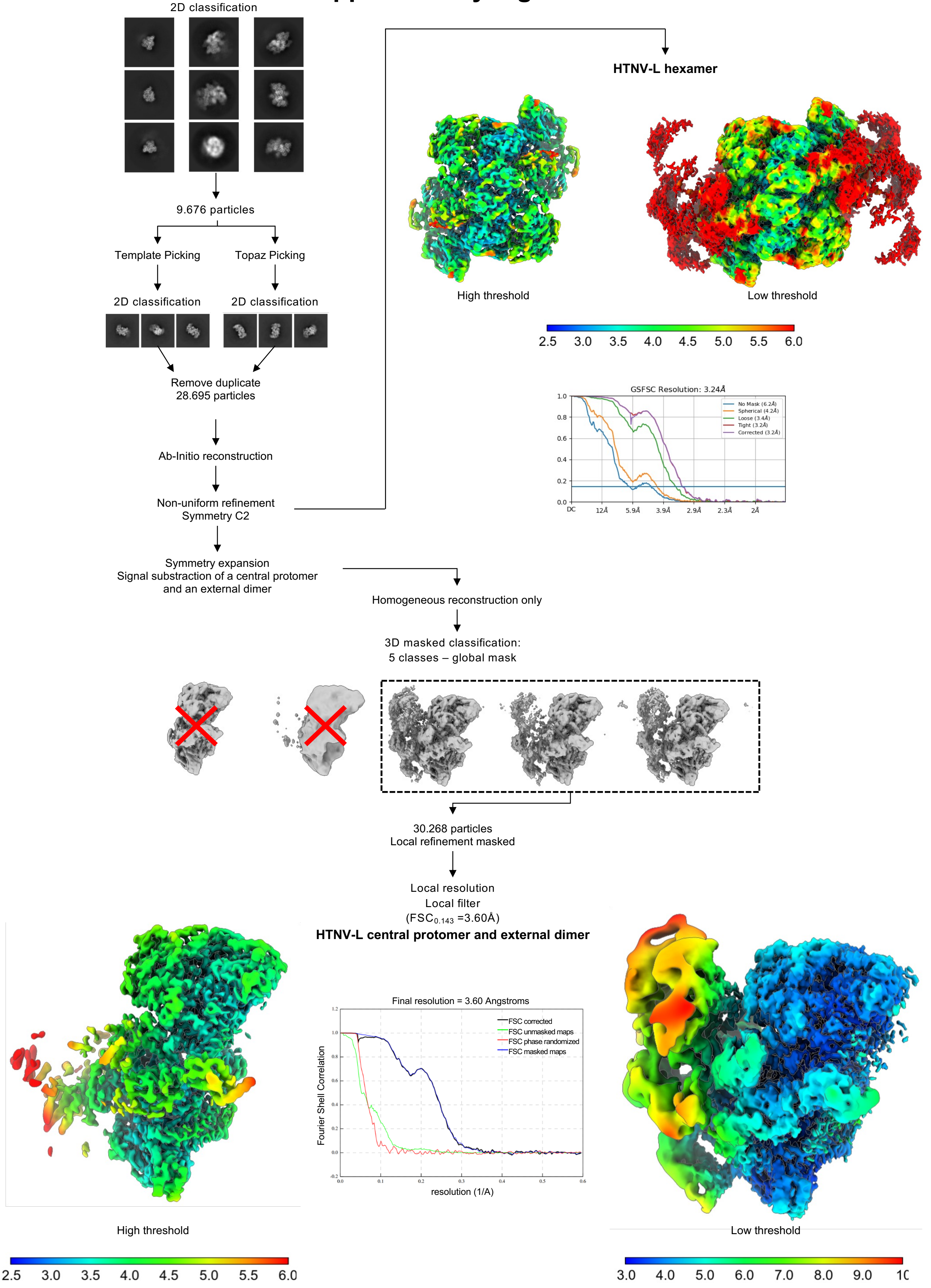


**Supplementary Figure 3 Image processing strategy to obtain hexameric apo HTNV-L**

Image processing strategy used to obtain HTNV-L hexamer. The strategy is displayed from the 1st round of 2D class averages also shown in **Supplementary Figure 2.** 2D class averages, 3D class averages and the final reconstructions are displayed. Region used for masking are indicated with a dotted line. 3D class averages chosen for further processing are surrounded by a rectangle dotted line. Electron density maps are colored according to the local resolution. Fourier Shell Correlation curves (FSC) are displayed.

### Supplementary Figure 4

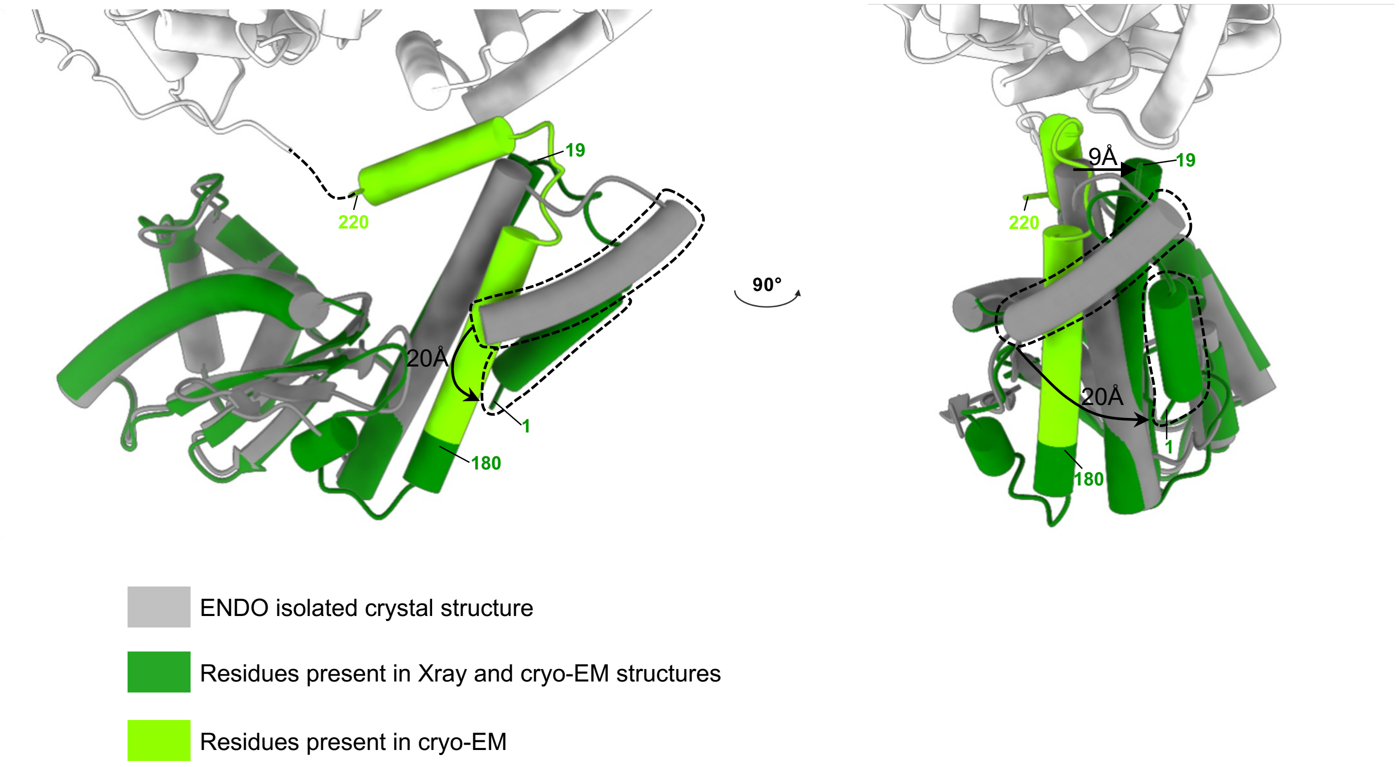


**Supplementary Figure 4 Comparison of the isolated ENDO structure from X-ray crystallography and the ENDO from the full-length HTNV-L**

Superposition of the isolated ENDO structure from X-ray crystallography colored in gray and the ENDO from the full-length HTNV-L colored in green. In the cartoon representation, residues in dark green are the ones present in both structures, and residues in light green are the ones present only in the cryo-EM full-length structure. The rotation amplitude is indicated. The $\alpha$-helix comprising residues 1 to 18 that moves significantly is surrounded by a dotted line.

### Supplementary Figure 5

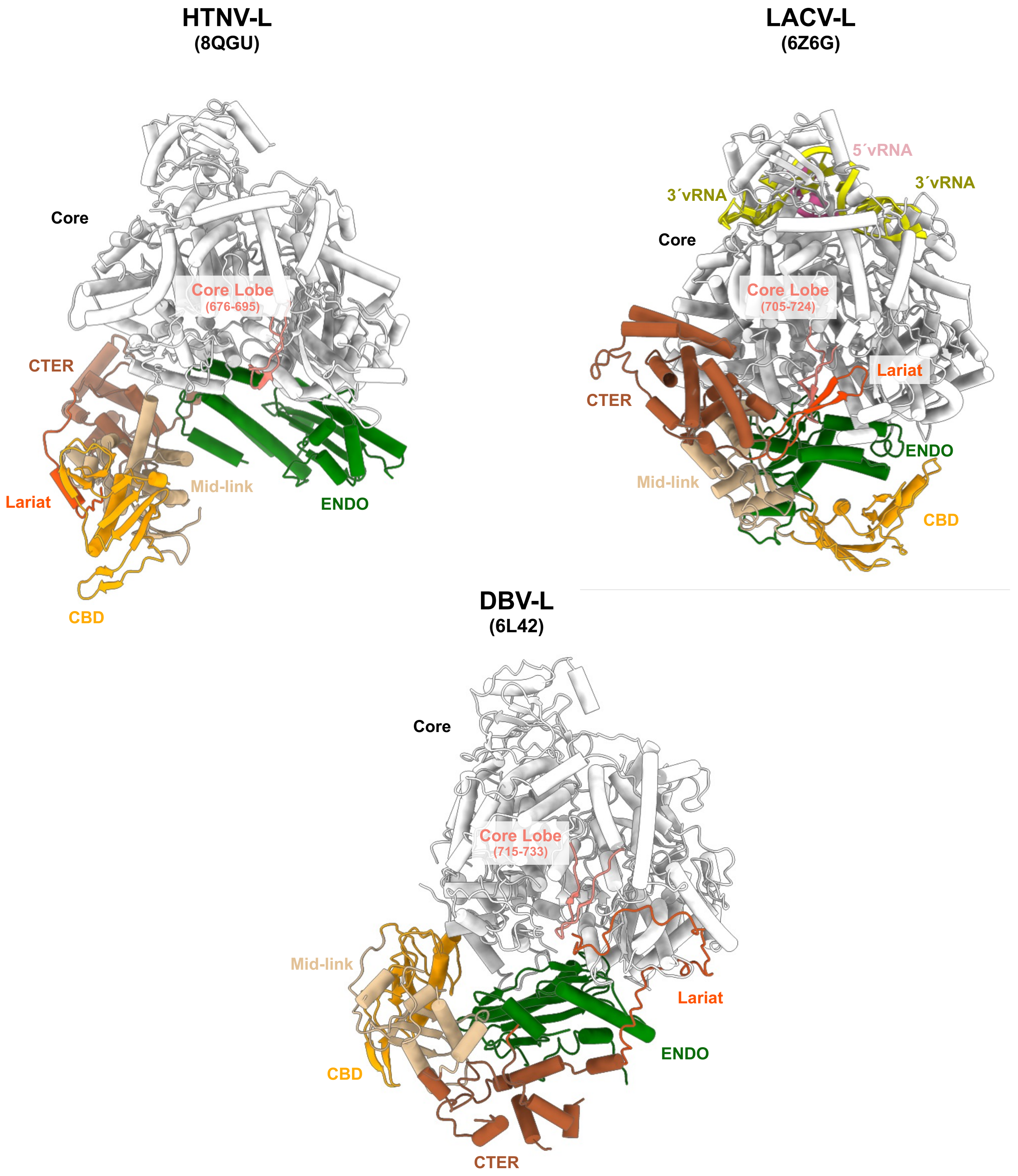


**Supplementary Figure 5 Comparison of lariat protrusion in Bunyaviruses**

HTNV-L, LACV-L and DBV-L are shown as cartoon and colored as in **Fig.2**. The β-hairpin strut of LACV-L and the lariat of HTNV-L and DBV-L are displayed in red. The β-hairpin present on the core-lobe of the three polymerases is colored in salmon.

### Supplementary Figure 6

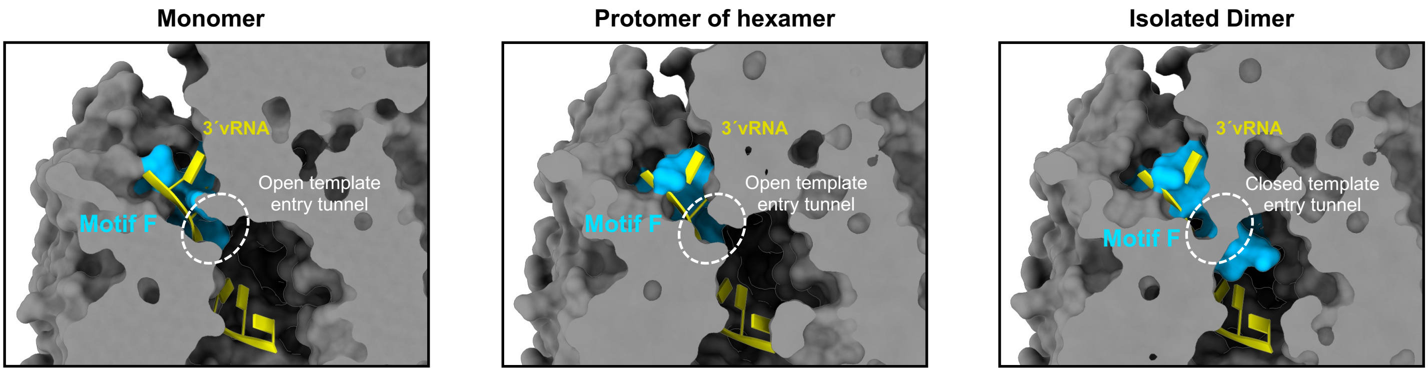


**Supplementary Figure 6 Motif F location compared to the template entry tunnel**

HTNV-L cores from HTNV-L apo monomer (left), dimer (middle) and hexamer (right) are colored as a white surface and clipped to visualize the 3´vRNA template entry tunnel. The location of the motif F is shown as a blue surface. The 3′vRNA end that originates from HTNV-L in pre-initiation (PDB 8C4U) is displayed as yellow cartoon.

### Supplementary Figure 7

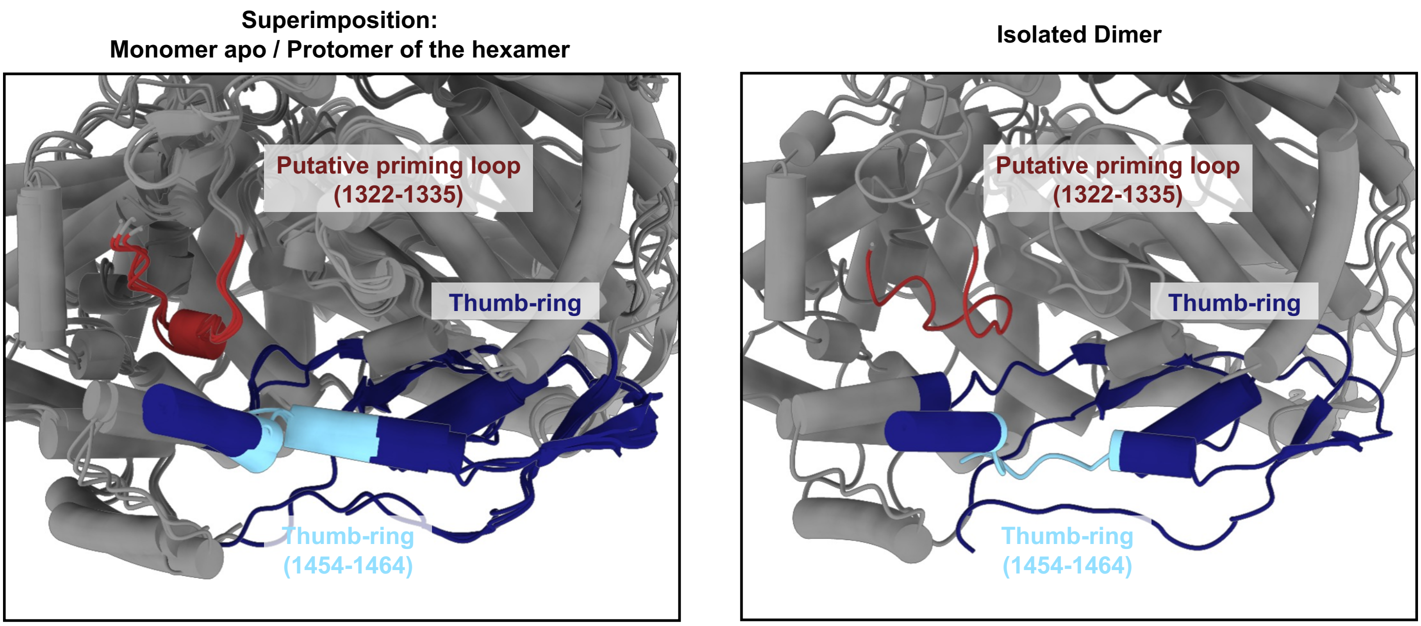


**Supplementary Figure 7 Comparison of the position of the putative priming loop in the different oligomers**

Zoom on the putative priming loop (or template exit plug) colored in dark red and thumb-ring colored in dark blue. The putative priming loop and thumb-ring region 1454-1464, that is colored in light blue, change their organization in the isolated HTNV-L dimer (right panel) compared to HTNV-L monomer and hexamer (left panel).

#### Supplementary Figure 8

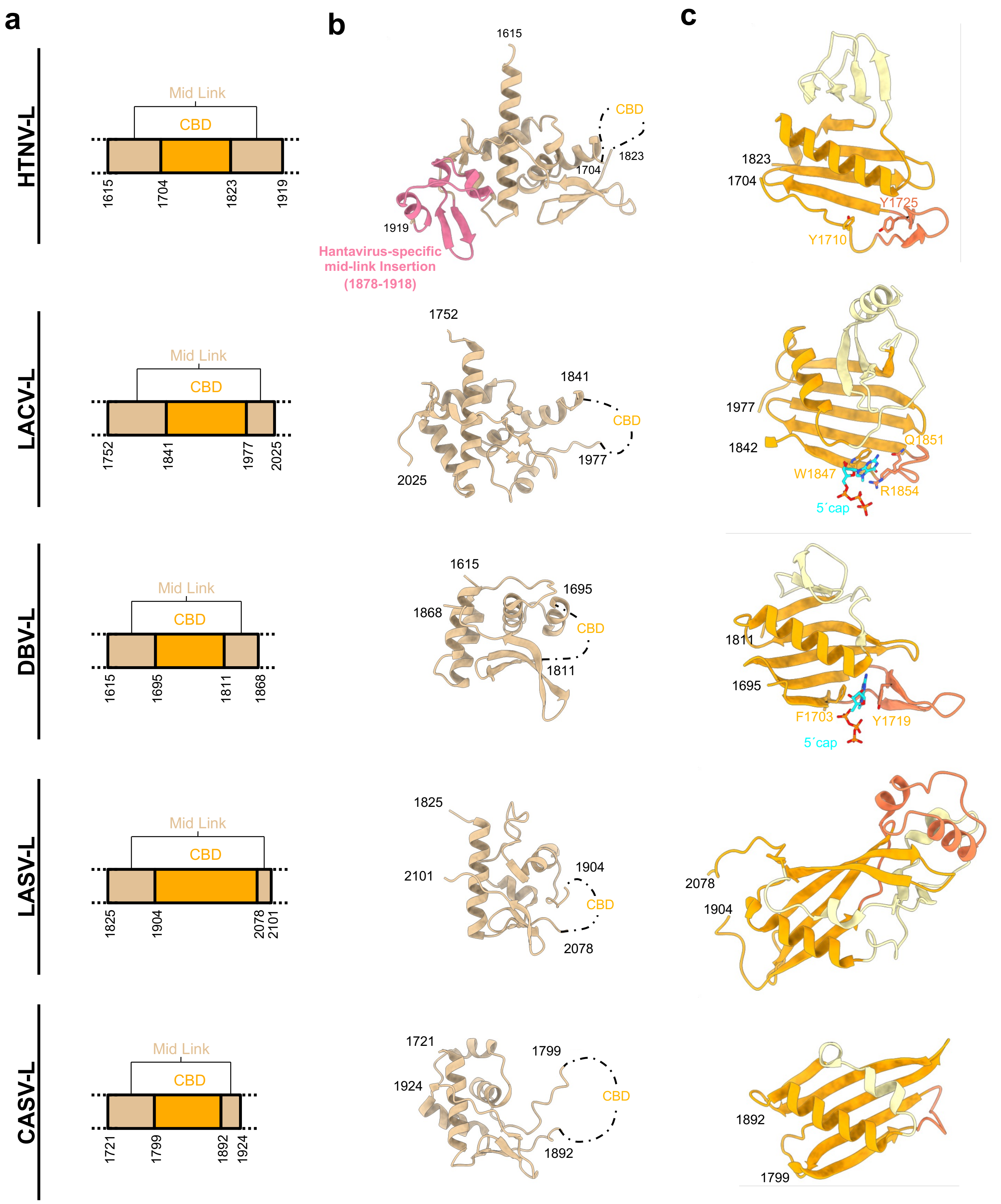


**Supplementary Figure 8 Mid-link and CBD organization in Bunyaviruses**

**a** Schematic representation of HTNV-L domain structure zoomed on the mid-link and the CBD.

**b** Mid-link domains from different Bunyaviruses colored in beige. The hantavirus-specific insertion is colored in light-pink.

**c** CBD domains from different sNSV. The central β-sheet and α-helix that are common to all Bunyavirus CBD are shown in orange cartoon. The β-hairpin insertion that is likely to be essential for cap binding is shown in dark orange. The region specific to each bunyavirus family is shown in light yellow. For LACV-L and DBV-L, the cap is shown as blue stick and residues that stack the cap are displayed as orange sticks. For HTNV-L and LASV-L, the residues that are likely to stack the cap due to their positioning are shown as sticks and labeled. CASV-L is missing the β-hairpin insertion and cannot stack the cap.

### Supplementary Figure 9

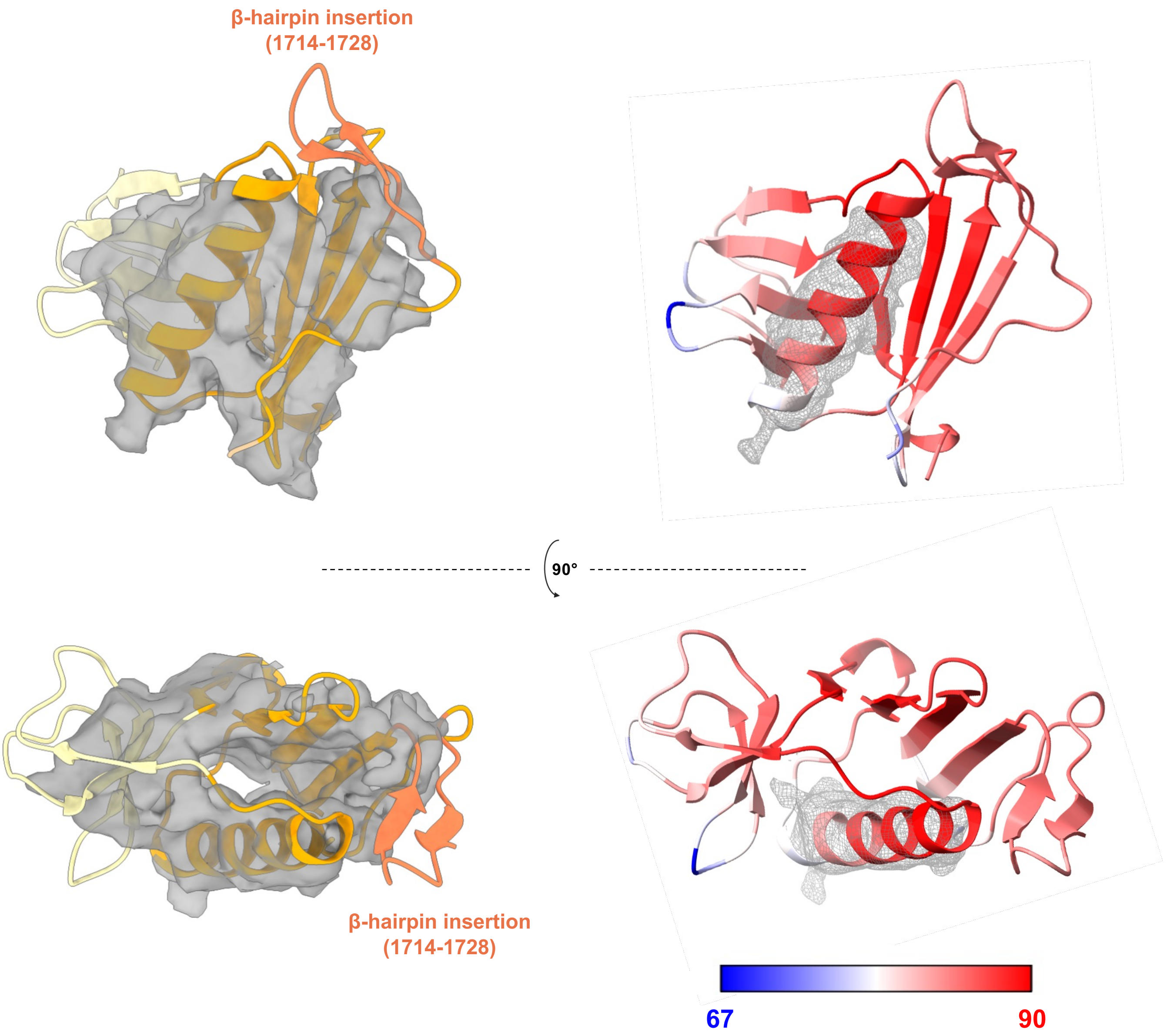


**Supplementary Figure 9 Cryo-EM envelope of the CBD**

On the left, the HTNV-L CBD model is shown as cartoon and colored as in **Fig. 2.** The corresponding cryo-EM density is shown as transparent gray surface. Although the resolution of this part remains low, the global fit is unambiguous as the global shape corresponds, with the central β-sheet and α-helix and the hantavirus-specific insertion that can be positioned in density. The β-hairpin insertion is not visible due to a too large flexibility.

On the right, the model is colored according to Alphafold pLDDT criteria. The electron density that corresponds to the α-helix is shown at a different threshold to show that it enables an unambiguous overall positioning of the CBD.

### Supplementary Figure 10

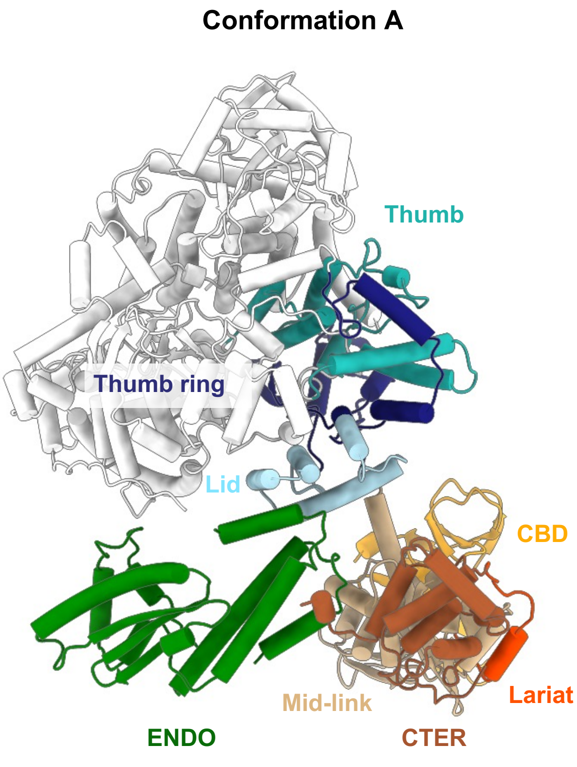


**Supplementary Figure 10 Interaction of the ENDO and the C-terminal domains with the polymerase cores in the conformers A and B**

Polymerase cores of HTNV-L conformers A and B are displayed as white cartoon. The domains that interact with either the ENDO or the C-terminal domains are colored: the lid in light blue, the thumb in light sea green and the thumb-ring in midnight blue.

### Supplementary Figure 11

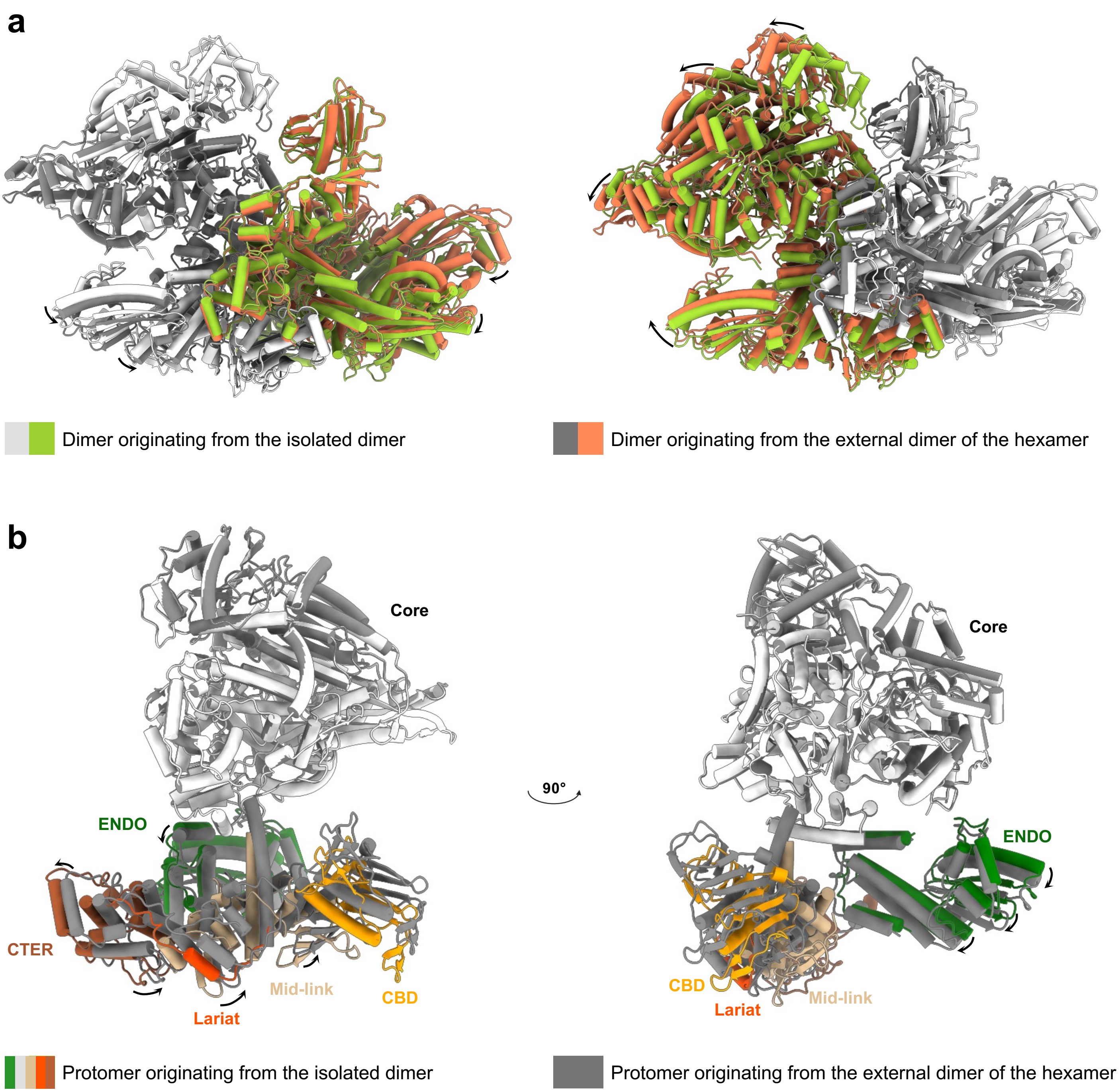


**Supplementary Figure 11 Comparison of dimers A in the isolated dimer and in the external dimer of the hexamer**

**a** superimposition of HTNV-L dimers A originating from the isolated dimer (colored in white and green) and from the external dimer of the hexamer (colored in grey and coral). A protomer originating from each dimer is superimposed and the relative rotation of the second protomer is indicated.

**b** superimposition of HTNV-L protomer originating from the isolated dimer and from the external dimer of the hexamer, with their core respectively colored in white and grey. The small rotations of the ENDO and the C-terminal domain, that are rigidly fitted in the density, are indicated.

### Supplementary Figure 12

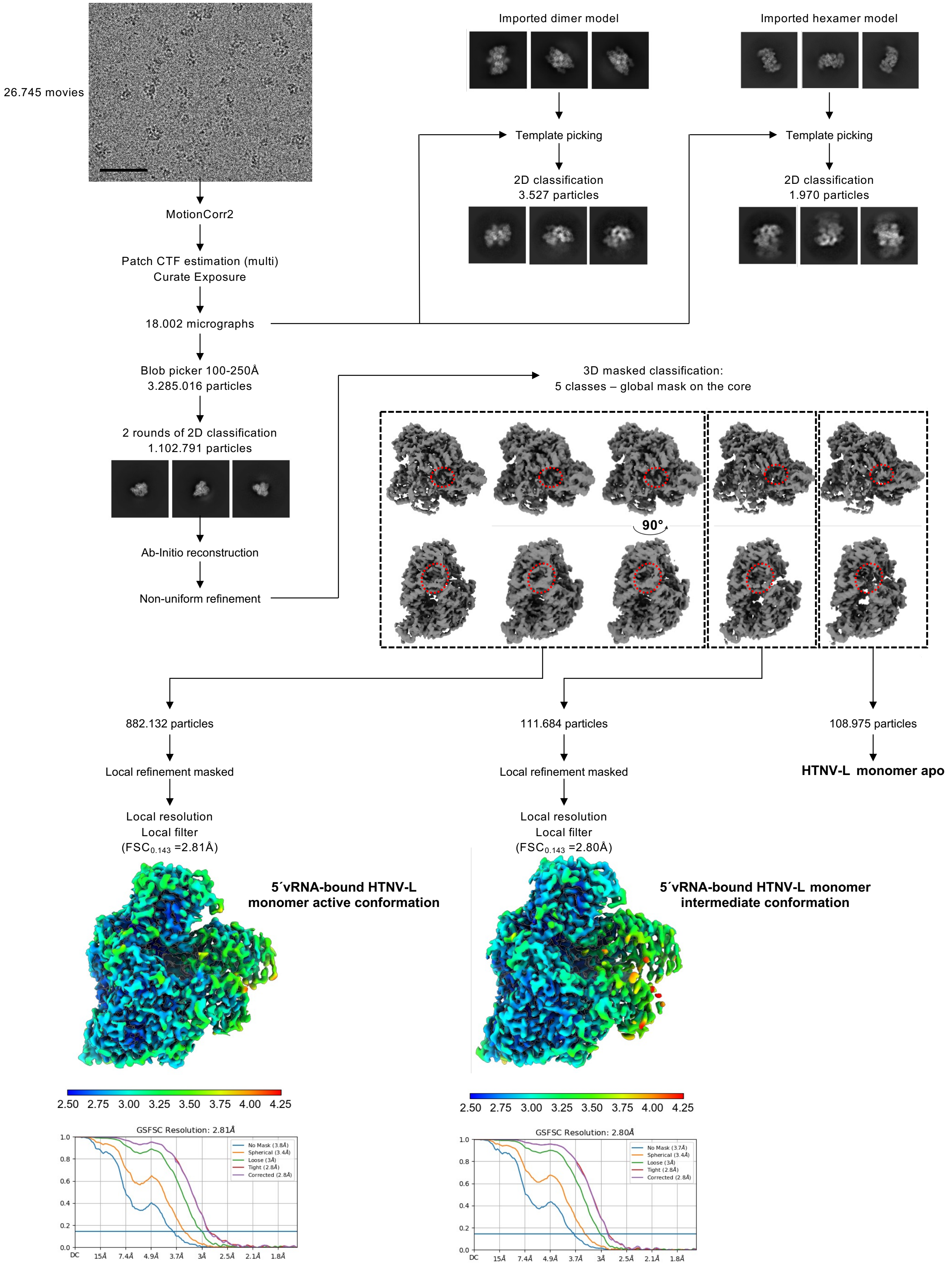


**Supplementary Figure 12 Image processing strategy to obtain monomeric 5′vRNA-bound HTNV-L structures**

A representative image of 5′vRNA-bound HTNV-L is displayed. The scale bar corresponds to 50 nm. Picking on the dataset did not reveal any dimers or hexamers. Import of dimers and hexamers of the apo HTNV-L dataset followed by template picking selected only a very low number of dimers and hexamers, preventing further processing.

The image processing workflow including 2D class averages, 3D class averages and the final reconstructions of monomeric 5′vRNA-bound HTNV-L are displayed. Electron density maps are colored according to the local resolution. Fourier Shell Correlation curves (FSC) are displayed.

### Supplementary Figure 13

**Superimposition:**
**Apo / 5´vRNA bound intermediate conformation**

**Superimposition:**
**Apo / 5´vRNA  bound active conformation**

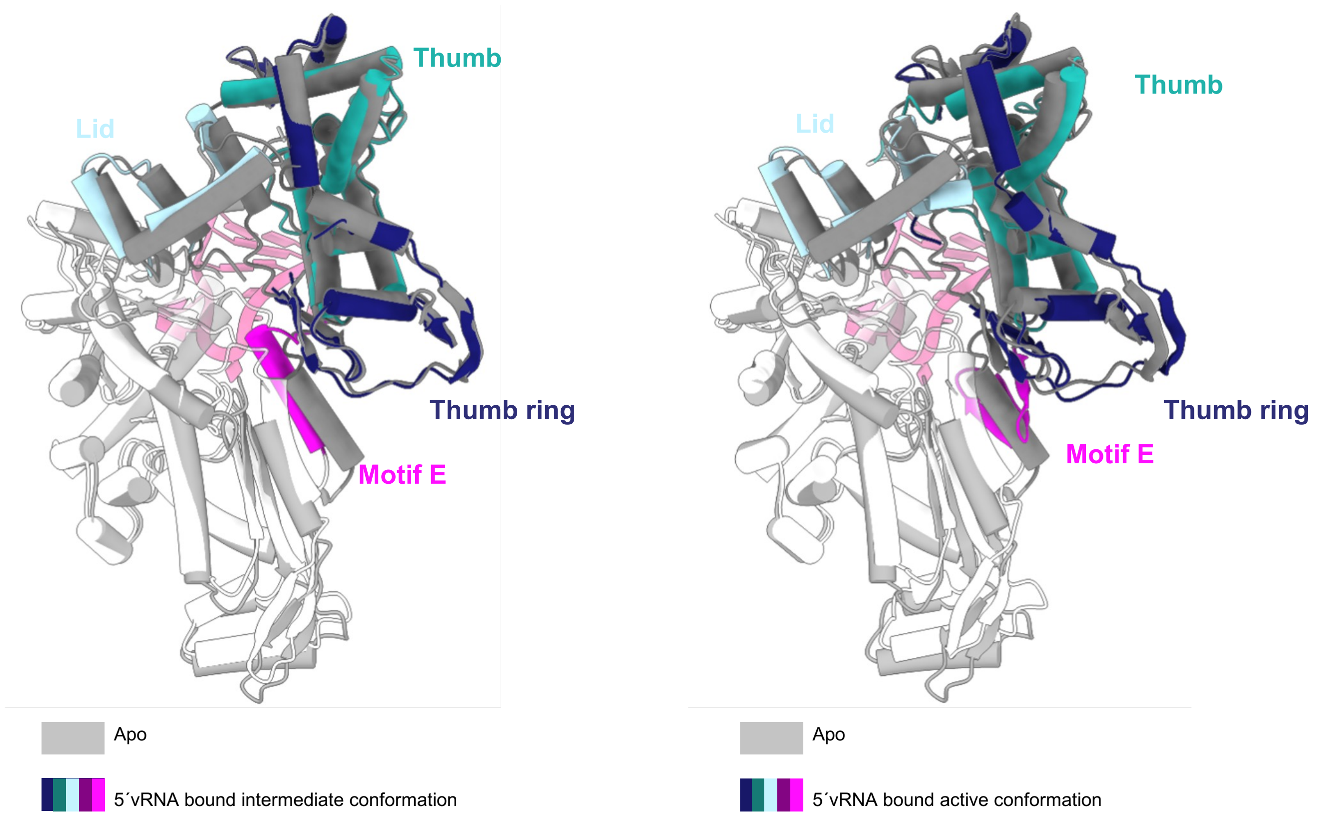


**Supplementary Figure 13 Comparison of HTNV-L polymerase cores**

HTNV-L apo monomeric structure is used as reference for superimposition and is displayed in white. It is superimposed with 5′vRNA-bound HTNV-L intermediate conformation (left) and 5′vRNA-bound active conformation (right). They are displayed as cartoon and colored in grey excepting their thumb, thumb-ring, lid and motif E that are respectively colored in light sea green, midnight blue, light blue and magenta.

### Supplementary Figure 14

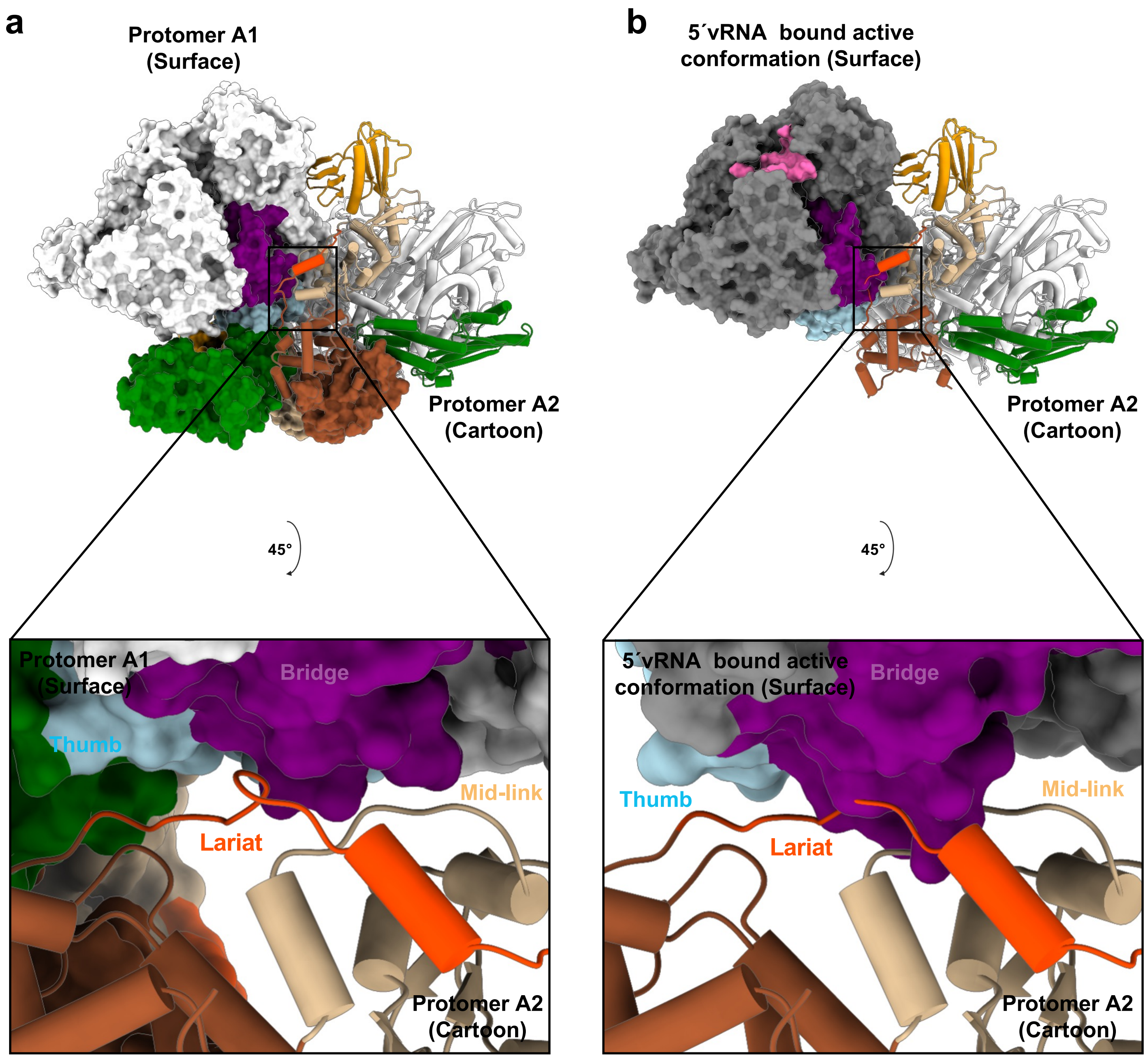


**Supplementary Figure 14 Superimposition the 5'vRNA-bound monomeric HTNV-L structure onto the HTNV-L isolated dimeric structure reveals clashes.**

**a** Structure of HTNV-L isolated dimer with protomer 1 shown as surface and protomer 2 shown as cartoon. The domains are colored as in **Fig.2** except the bridge and the thumb of protomer 1 that are respectively displayed in purple and in light blue. The interaction between the lariat and the bridge is visible on the zoom displayed at the bottom.

**b** When superimposing the 5'vRNA-bound HTNV-L core structure on protomer 1, clashes appear with protomer 2. Protomer 1 is not shown, protomer 2 is shown as in **a**, 5'vRNA-bound HTNV-L core is shown as a grey surface with the bridge and the thumb domains colored in purple and in light blue. The zoom at the bottom identifies clashes between the lariat of protomer 2 and the bridge of protomer 1.

### Supplementary Figure 15

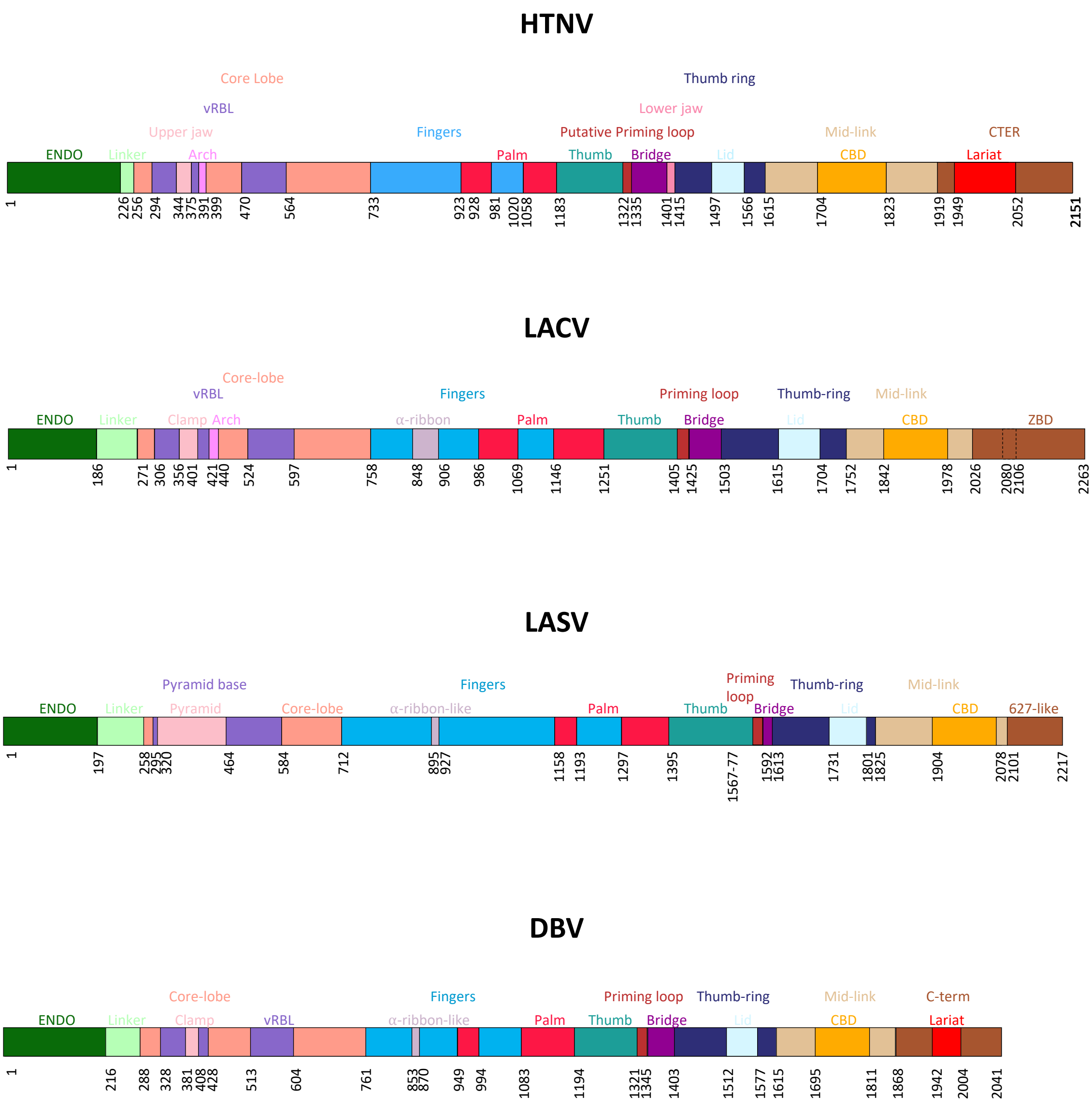


**Supplementary Figure 15 Schematic representation of Bunyaviruses domain structure**

Each domain is colored and its extreme residues are numbered.

### Supplementary Figure 16

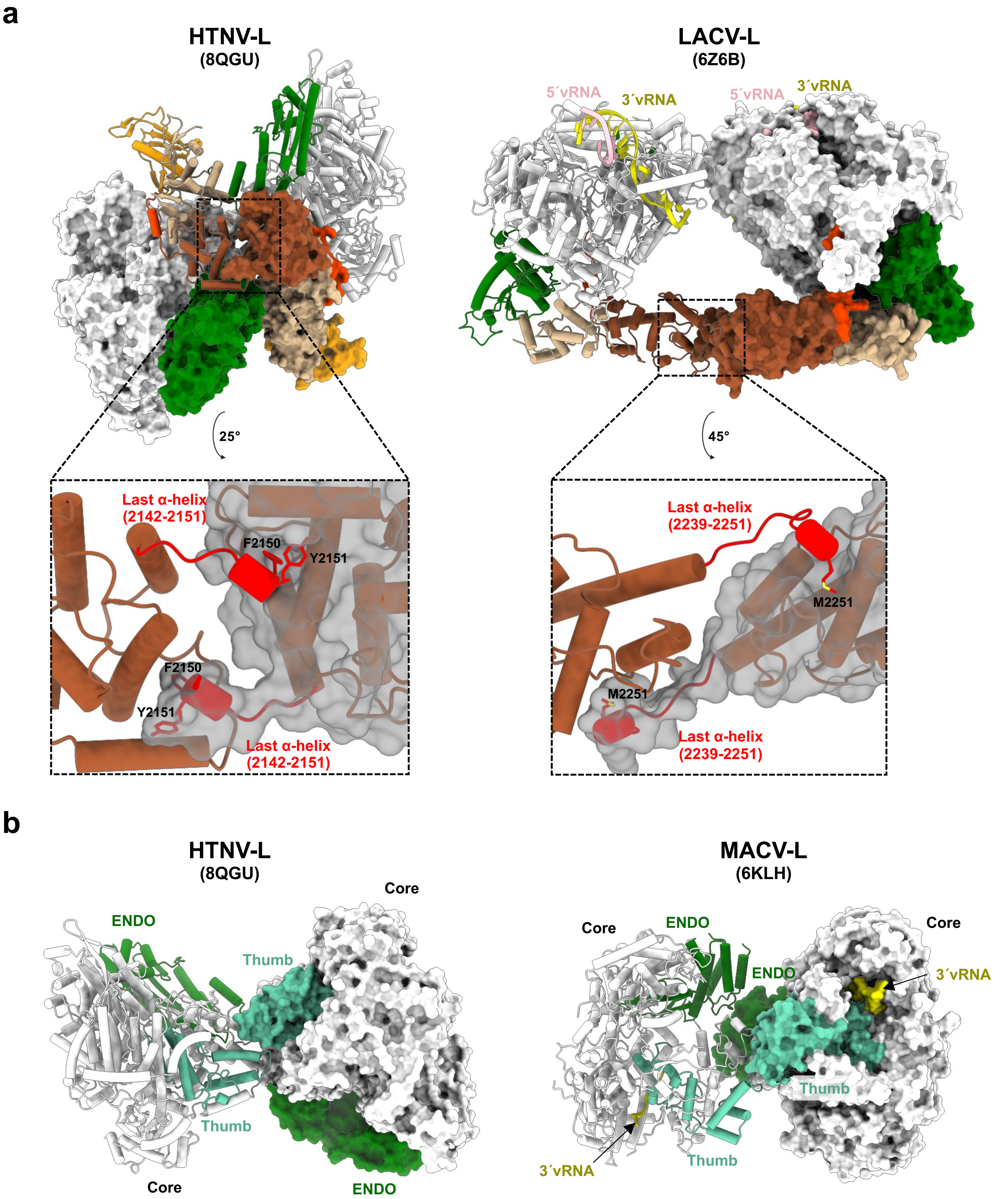


**Supplementary Figure 16 Comparison of HTNV-L dimer with other sNSV polymerase dimers**

**a** HTNV-L and LACV-L dimers colored as in **Fig. 3**. One protomer is shown as surface and the other as cartoon. LACV-L is bound to 5′ and 3′vRNA ends that are respectively shown as light pink and yellow cartoons. At the bottom, zoom on the C-terminal ends, with the surface of one protomer shown. The C-terminal extremities that swap in the other protomer are shown in red, with the conserved hydrophobic C-terminal residues shown as sticks.

**b** HTNV-L and MACV-L dimers with one protomer shown as cartoon and the second displayed as a surface. The cores are shown in white with the exception of the thumbs that are colored in light sea green. The ENDO are colored in forest green. The 3′vRNA that is bound to MACV-L dimer is shown in yellow.

#### SUPPLEMENTARY FIGURE LEGENDS

**Supplementary Figure 1 HTNV-L purification and characterization of its oligomerization**

**a** Size-exclusion chromatography profile on a superdex 200 increase 3.2/300 column. The optical density (OD) for 255 nm and 280 nm is indicated respectively in red and green as a function of the elution volume. The three fractions analyzed in **b** and **c** are indicated with arrows.

**b** SDS-PAGE gels of the 3 fractions indicated in **a**. The molecular weight marker (MW) is shown on the left.

**c** Mass photometry of the 3 fractions indicated in **a**. The mass in kDa and the percentage of each specie detected is indicated.

**Supplementary Figure 2 Image processing strategy to obtain monomeric and dimeric apo and HTNV-L**

A representative image of HTNV-L apo is displayed. The scale bar corresponds to 50 nm. The image processing workflow including 2D class averages, 3D class averages and the final reconstructions are displayed. Regions used for masking are indicated with a dotted line. 3D class averages chosen for further processing are surrounded by a rectangle dotted line. Electron density maps are colored according to the local resolution. Gold-standard Fourier Shell Correlation curves (FSC) are displayed.

**Supplementary Figure 3 Image processing strategy to obtain hexameric apo HTNV-L**

Image processing strategy used to obtain HTNV-L hexamer. The strategy is displayed from the 1$^{st}$ round of 2D class averages also shown in **Supplementary Figure 2.** 2D class averages, 3D class averages and the final reconstructions are displayed. Region used for masking are indicated with a dotted line. 3D class averages chosen for further processing are surrounded by a rectangle dotted line. Electron density maps are colored according to the local resolution. Fourier Shell Correlation curves (FSC) are displayed.

**Supplementary Figure 4 Comparison of the isolated ENDO structure from X-ray crystallography and the ENDO from the full-length HTNV-L**

Superposition of the isolated ENDO structure from X-ray crystallography colored in gray and the ENDO from the full-length HTNV-L colored in green. In the cartoon representation, residues in dark green are the ones present in both structures, and residues in light green are the ones present only in the cryo-EM full-length structure. The rotation amplitude is indicated. The $\alpha$-helix comprising residues 1 to 18 that moves significantly is surrounded by a dotted line.

**Supplementary Figure 5 Comparison of lariat protrusion in Bunyaviruses**

HTNV-L, LACV-L and DBV-L are shown as cartoon and colored as in **Fig.2**. The $\beta$-hairpin strut of LACV-L and the lariat of HTNV-L and DBV-L are displayed in red. The $\beta$-hairpin present on the core-lobe of the three polymerases is colored in salmon.

**Supplementary Figure 6 Motif F location compared to the template entry tunnel**

HTNV-L cores from HTNV-L apo monomer (left), dimer (middle) and hexamer (right) are colored as a white surface and clipped to visualize the 3´vRNA template entry tunnel. The location of the motif F is shown as a blue surface. The 3'vRNA end that originates from HTNV-L in pre-initiation (PDB 8C4U) is displayed as yellow cartoon.

**Supplementary Figure 7 Comparison of the position of the putative priming loop in the different oligomers**

Zoom on the putative priming loop (or template exit plug) colored in dark red and thumb-ring colored in dark blue. The putative priming loop and thumb-ring region 1454-1464, that is colored in light blue, change their organization in the isolated HTNV-L dimer (right panel) compared to HTNV-L monomer and hexamer (left panel).

**Supplementary Figure 8 Mid-link and CBD organization in Bunyaviruses**

**a** Schematic representation of HTNV-L domain structure zoomed on the mid-link and the CBD.

**b** Mid-link domains from different Bunyaviruses colored in beige. The hantavirus-specific insertion is colored in light-pink.

**c** CBD domains from different sNSV. The central $\beta$-sheet and $\alpha$-helix that are common to all Bunyavirus CBD are shown in orange cartoon. The $\beta$-hairpin insertion that is likely to be essential for cap binding is shown in dark orange. The region specific to each bunyavirus family is shown in light yellow. For LACV-L and DBV-L, the cap is shown as blue stick and residues that stack the cap are displayed as orange sticks. For HTNV-L and LASV-L, the residues that are likely to stack the cap due to their positioning are shown as sticks and labeled. CASV-L is missing the β-hairpin insertion and cannot stack the cap.

**Supplementary Figure 9 Cryo-EM envelope of the CBD**

On the left, the HTNV-L CBD model is shown as cartoon and colored as in **Fig. 2.** The corresponding cryo-EM density is shown as transparent gray surface. Although the resolution of this part remains low, the global fit is unambiguous as the global shape corresponds, with the central β-sheet and α-helix and the hantavirus-specific insertion that can be positioned in density. The β-hairpin insertion is not visible due to a too large flexibility.

On the right, the model is colored according to Alphafold pLDDT criteria. The electron density that corresponds to the α-helix is shown at a different threshold to show that it enables an unambiguous overall positioning of the CBD.

**Supplementary Figure 10 Interaction of the ENDO and the C-terminal domains with the polymerase cores in the conformers A and B**

Polymerase cores of HTNV-L conformers A and B are displayed as white cartoon. The domains that interact with either the ENDO or the C-terminal domains are colored: the lid in light blue, the thumb in light sea green and the thumb-ring in midnight blue.

**Supplementary Figure 11 Comparison of dimers A in the isolated dimer and in the external dimer of the hexamer**

**a** superimposition of HTNV-L dimers A originating from the isolated dimer (colored in white and green) and from the external dimer of the hexamer (colored in grey and coral). A protomer originating from each dimer is superimposed and the relative rotation of the second protomer is indicated.

**b** superimposition of HTNV-L protomer originating from the isolated dimer and from the external dimer of the hexamer, with their core respectively colored in white and grey. The small rotations of the ENDO and the C-terminal domain, that are rigidly fitted in the density, are indicated.

**Supplementary Figure 12 Image processing strategy to obtain monomeric 5’vRNA-bound HTNV-L structures**

A representative image of 5’vRNA-bound HTNV-L is displayed. The scale bar corresponds to 50 nm. Picking on the dataset did not reveal any dimers or hexamers. Import of dimers and hexamers of the apo HTNV-L dataset followed by template picking selected only a very low number of dimers and hexamers, preventing further processing.

The image processing workflow including 2D class averages, 3D class averages and the final reconstructions of monomeric 5’vRNA-bound HTNV-L are displayed. Electron density maps are colored according to the local resolution. Fourier Shell Correlation curves (FSC) are displayed.

**Supplementary Figure 13 Comparison of HTNV-L polymerase cores**

HTNV-L apo monomeric structure is used as reference for superimposition and is displayed in white. It is superimposed with 5’vRNA-bound HTNV-L intermediate conformation (left) and 5’vRNA-bound active conformation (right). They are displayed as cartoon and colored in grey excepting their thumb, thumb-ring, lid and motif E that are respectively colored in light sea green, midnight blue, light blue and magenta.

**Supplementary Figure 14 Superimposition the 5’vRNA-bound monomeric HTNV-L structure onto the HTNV-L isolated dimeric structure reveals clashes.**

**a** Structure of HTNV-L isolated dimer with protomer 1 shown as surface and protomer 2 shown as cartoon. The domains are colored as in Fig.2 except the bridge and the thumb of protomer 1 that are respectively displayed in purple and in light blue. The interaction between the lariat and the bridge is visible on the zoom displayed at the bottom.

**b** When superimposing the 5’vRNA-bound HTNV-L core structure on protomer 1, clashes appear with protomer 2. Protomer 1 is not shown, protomer 2 is shown as in **a**, 5’vRNA-bound HTNV-L core is shown as a grey surface with the bridge and the thumb domains colored in purple and in light blue. The zoom at the bottom identifies clashes between the lariat of protomer 2 and the bridge of protomer 1.

**Supplementary Figure 15 Schematic representation of Bunyaviruses domain structure**

Each domain is colored and its extreme residues are numbered.

**Supplementary Figure 16 Comparison of HTNV-L dimer with other sNSV polymerase dimers**

**a** HTNV-L and LACV-L dimers colored as in **Fig. 4**. One protomer is shown as surface and the other as cartoon. LACV-L is bound to 5′ and 3′vRNA ends that are respectively shown as light pink and yellow cartoons. At the bottom, zoom on the C-terminal ends, with the surface of one protomer shown. The C-terminal extremities that swap in the other protomer are shown in red, with the conserved hydrophobic C-terminal residues shown as sticks.

**b** HTNV-L and MACV-L dimers with one protomer shown as cartoon and the second displayed as a surface. The cores are shown in white with the exception of the thumbs that are colored in light sea green. The ENDO are colored in forest green. The 3′vRNA that is bound to MACV-L dimer is shown in yellow.
