## Supplementary figures and images for "Structural characterization of the oligomerization of full-length Hantaan virus polymerase into symmetric dimers and hexamers"

### Supplementary Data 1

#### Supplementary Data (1/3)

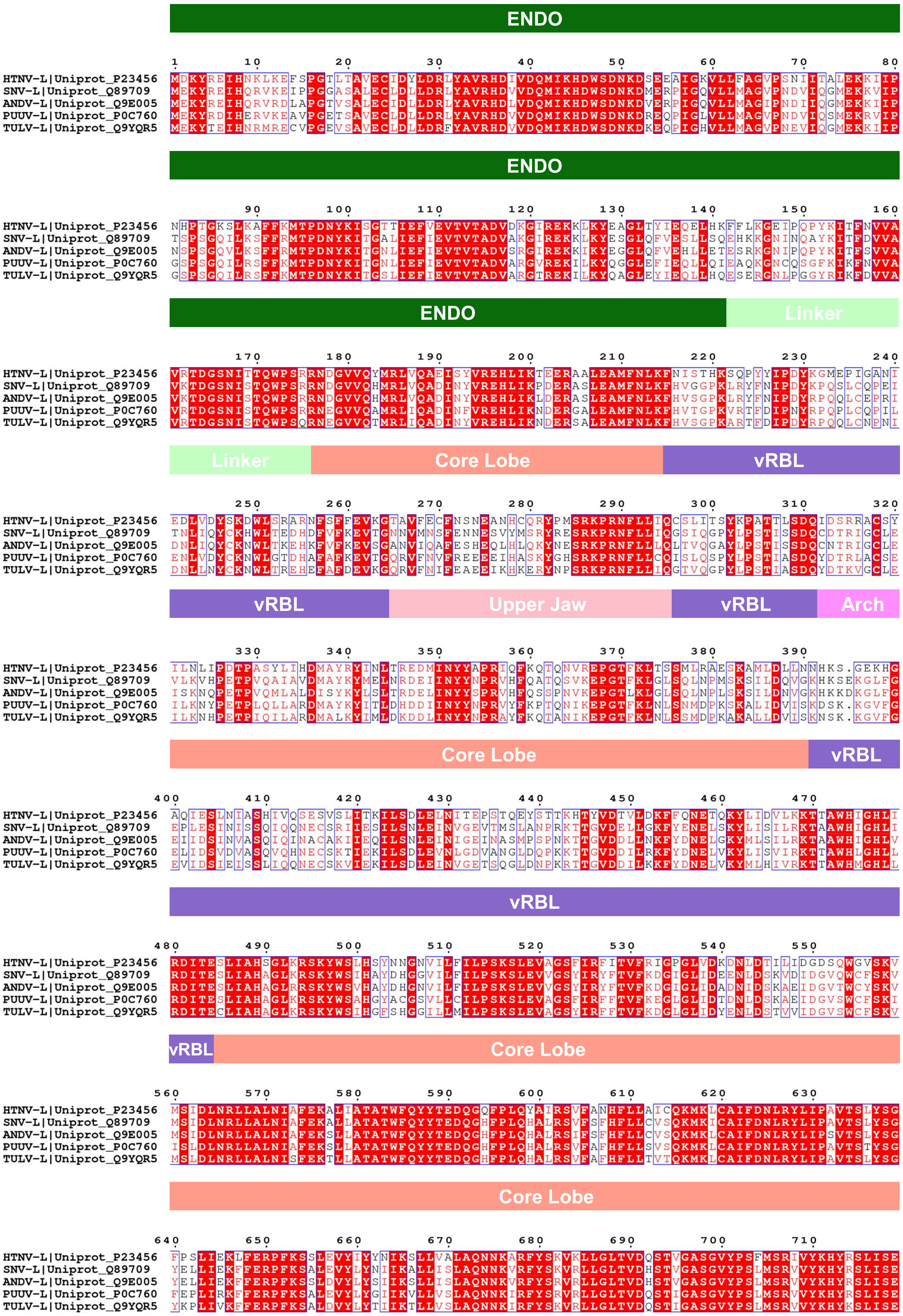

#### Supplementary Data (2/3)

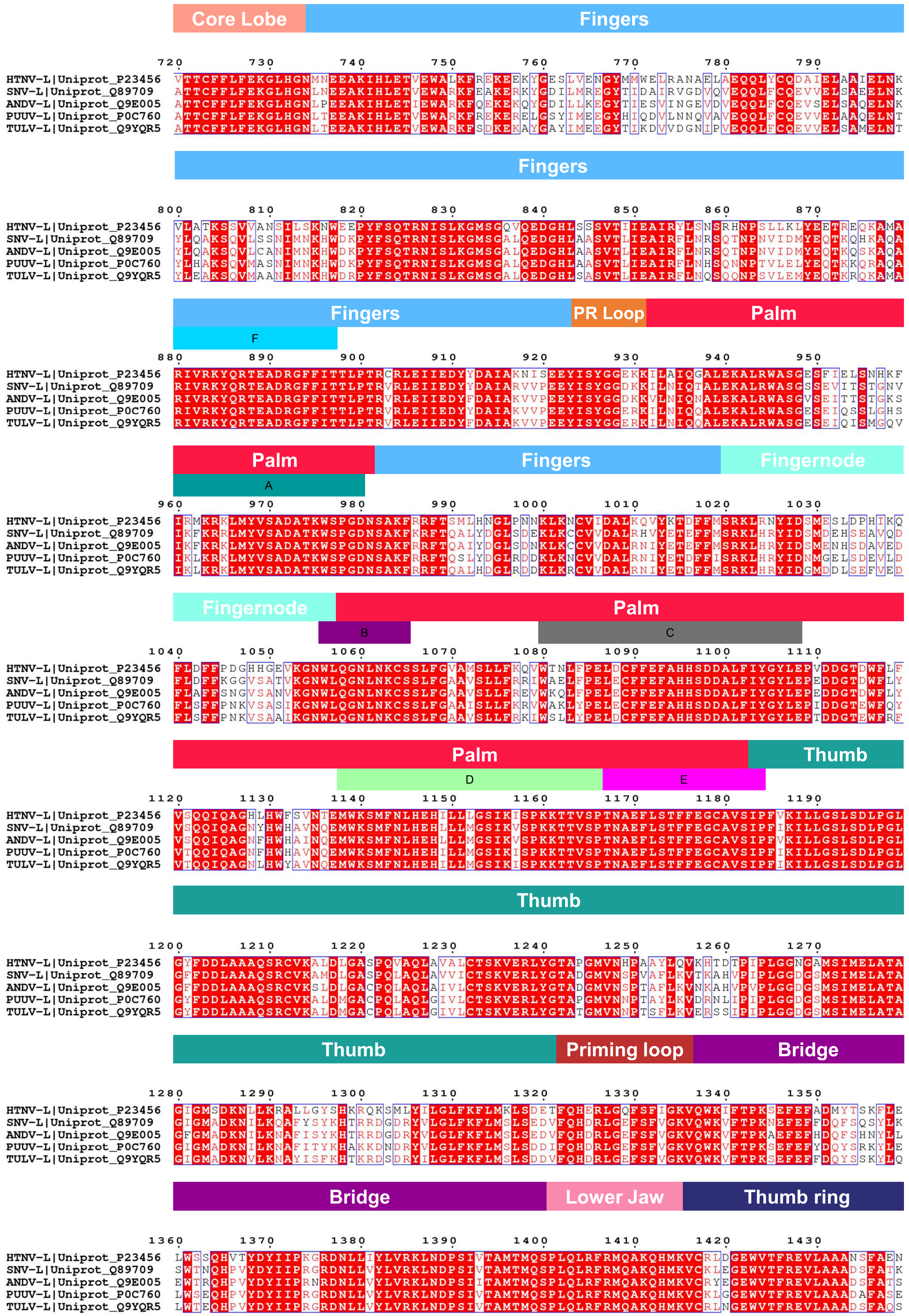

#### Supplementary Data (3/3)

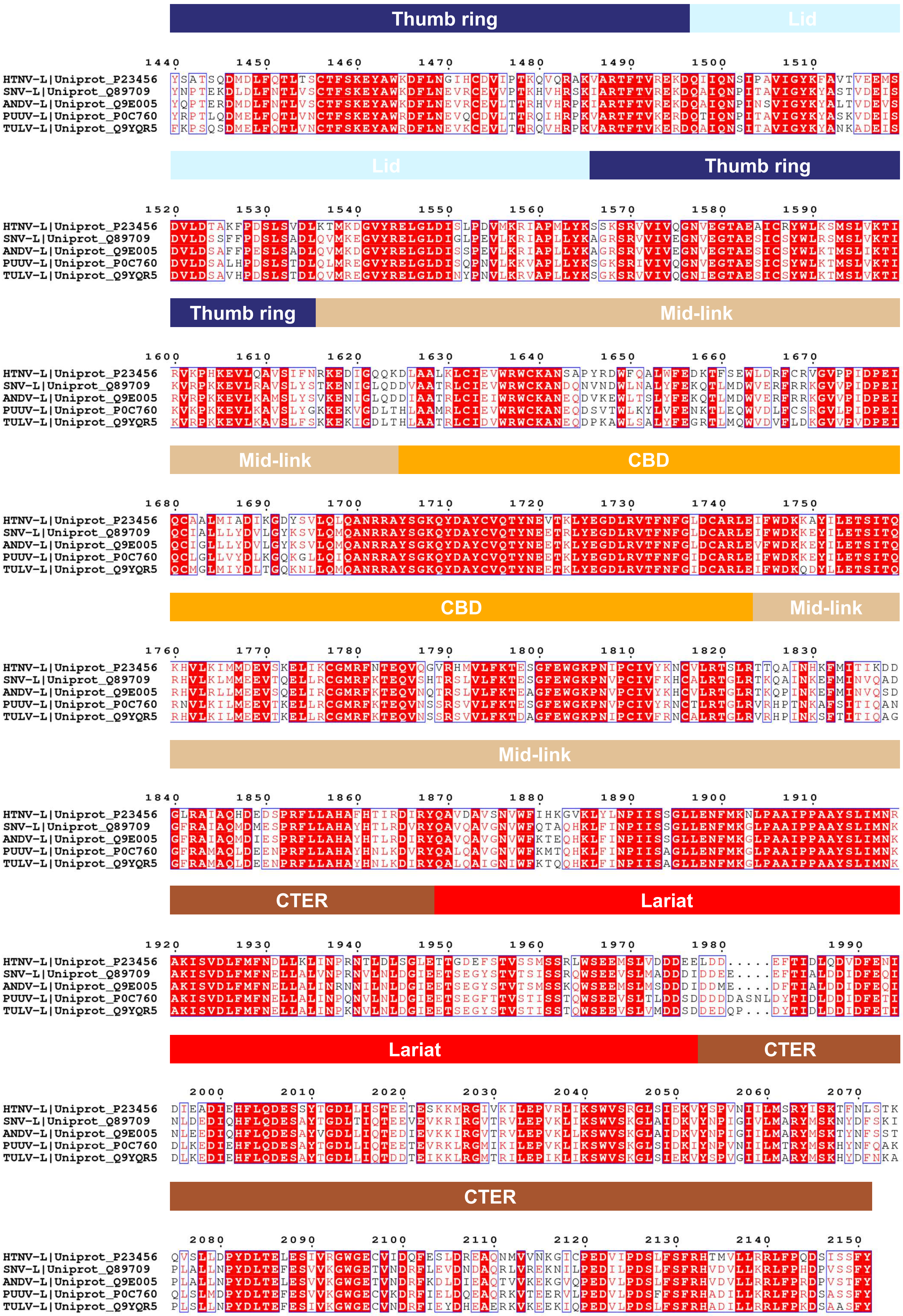
